# Chk2 sustains PLK1 activity in mitosis to ensure proper chromosome segregation

**DOI:** 10.1101/2024.03.08.584115

**Authors:** Elizabeth M. Black, Carlos Andrés Ramírez Parrado, Isabelle Trier, Wenxue Li, Yoon Ki Joo, Jennifer Pichurin, Yansheng Liu, Lilian Kabeche

## Abstract

Polo-like kinase 1 (PLK1) protects against genome instability by ensuring timely and accurate mitotic cell division. PLK1 activity is tightly regulated throughout the cell cycle. Although the pathways that initially activate PLK1 in G2 are well-characterized, the factors that directly regulate PLK1 in mitosis remain poorly understood. Here, we identify that human PLK1 activity is sustained by the DNA damage response kinase Checkpoint kinase 2 (Chk2) in mitosis. Chk2 directly phosphorylates PLK1 T210, a residue on its T-loop whose phosphorylation is essential for full PLK1 kinase activity. Loss of Chk2-dependent PLK1 activity causes increased mitotic errors, including chromosome misalignment, chromosome missegregation, and cytokinetic defects. Moreover, Chk2 deficiency increases sensitivity to PLK1 inhibitors, suggesting that Chk2 status may be an informative biomarker for PLK1 inhibitor efficacy. This work demonstrates that Chk2 sustains mitotic PLK1 activity and protects genome stability through discrete functions in interphase DNA damage repair and mitotic chromosome segregation.

## Introduction

Accurate mitotic progression is essential for proper cellular and organismal function, and errors in this process can both initiate and drive cancer development ^1–4^. One important protein that protects against such defects is Polo-like kinase 1 (PLK1), which promotes proper centrosome maturation, bipolar spindle formation, kinetochore-microtubule attachment, chromosome segregation, and cytokinesis ^5–12^. Tight spatial and temporal regulation of PLK1 is necessary to accomplish these pleiotropic functions ^13–17^, and even partial reduction in PLK1 activity can lead to genome instability, senescence, and failure to proliferate ^8, 18^. Thus, comprehensively identifying the factors that regulate mitotic PLK1 is important to better understand how mitotic cell division is controlled and identify novel pathways that protect genome stability.

PLK1 is initially activated in G2 through a highly synchronized positive feedback loop with the mitotic kinases CDK1 and Aurora A. Aurora A, in complex with its co-factor Bora, directly phosphorylates PLK1 T210, a residue on PLK1’s T-loop whose phosphorylation promotes PLK1 activity ^19–22^. Active PLK1 promotes CDK1 activity ^23–25^, which further supports Aurora A-Bora binding and activity as part of a bistable positive feedback loop that ensures PLK1, Aurora A, and CDK1 are highly active as they enter mitosis ^26, 27^. This positive feedback loop is interrupted in the presence of DNA damage when signaling proteins in the DNA damage response pathway inactivate CDK1 to trigger a reversible G2/M arrest ^28–30^.

Although the mechanisms that activate PLK1 in G2 are well-defined, we have an incomplete understanding of the factors that regulate PLK1 in mitosis. After mitotic entry, Bora is largely degraded, and Aurora A associates with other co-factors, including TPX2 ^20, 31–33^. While there is a small residual population of Aurora A-Bora that continues to sustain PLK1 activity in mitosis, its contribution is limited to only ∼20% of PLK1 activity ^34^. We know that PLK1 activity, however, must remain high throughout mitosis despite negative regulation from phosphatases ^35, 36^. This observation suggests that there may be previously undescribed factors that act on PLK1 to sustain its activity in mitosis.

Chk2 is an effector kinase in the DNA damage response pathway that has a well-characterized role to control cell cycle progression following DNA double-stranded breaks. In addition to this canonical role in the interphase DNA damage response, Chk2 has been implicated in a variety of mitotic processes, including checkpoint signaling, spindle formation, cytokinesis, and chromosome segregation, suggesting that Chk2 may have an important role in mitosis ^37–46^. These previous studies, however, utilized depletion methods such as knockdown or knockout that rely on natural protein turnover over multiple cell cycles, confounding the known interphase and putative mitotic functions of Chk2. These studies also lack a unifying mechanism to fully explain whether and how Chk2 could be executing these mitotic functions. Given that many of the defects associated with Chk2 deficiency mimic those observed following PLK1 inhibition ^8, 47^, we hypothesized that there may be a previously unreported relationship between Chk2 and PLK1. Here, we identify that Chk2 promotes mitotic PLK1 activity, and that inhibiting this pathway in mitosis causes chromosome segregation defects and genome instability. Our data suggest a model where this effect occurs in part through direct phosphorylation of PLK1 T210 by Chk2. We propose a new paradigm whereby the mechanisms that initially activate PLK1 are distinct from those that sustain it, and that Chk2 protects genome stability through discrete functions in interphase DNA damage response and mitotic chromosome segregation.

## Results

### Chk2 inhibition reduces mitotic PLK1 activity

To test whether Chk2 regulates mitotic PLK1 activity, we used nocodazole to arrest cells in mitosis, isolated mitotic cells via shake off, and subsequently treated cells with inhibitors of PLK1 (1μM BI-2536, PLK1i), Chk2 (10μM BLM-277, Chk2i), or a vehicle control (DMSO), and measured the abundance of phosphorylated TCTP Serine 46 (p-TCTP), a marker of PLK1 activity ^48^ (Fig. 1A, Fig. S1A). We observed that treatment with Chk2i caused a ∼25% decrease in p-TCTP in HeLa (cervical adenocarcinoma), RPE-1 (non-transformed retinal pigmented epithelium), PANC1 (pancreatic ductal adenocarcinoma), U2OS (osteosarcoma), and A549 (lung adenocarcinoma) cells. This effect is not due to degradation of PLK1 or TCTP, as treatment with inhibitors did not cause changes in total protein abundance (Fig. 1A, S1B-C). Inhibitor treatment also did not affect the abundance or foci count of γH2AX, a marker of DNA damage (Fig. S2) ^49^. p-TCTP was similarly affected following mitotic arrest with the Eg5 inhibitor S-trityl-L-cysteine (STLC, 10μM) and inhibitor treatment (Fig. 1B). Taken together, these data suggest that Chk2 kinase activity is necessary to sustain full PLK1 activity in mitosis in cancerous and noncancerous cell types.

**Fig. 1.**
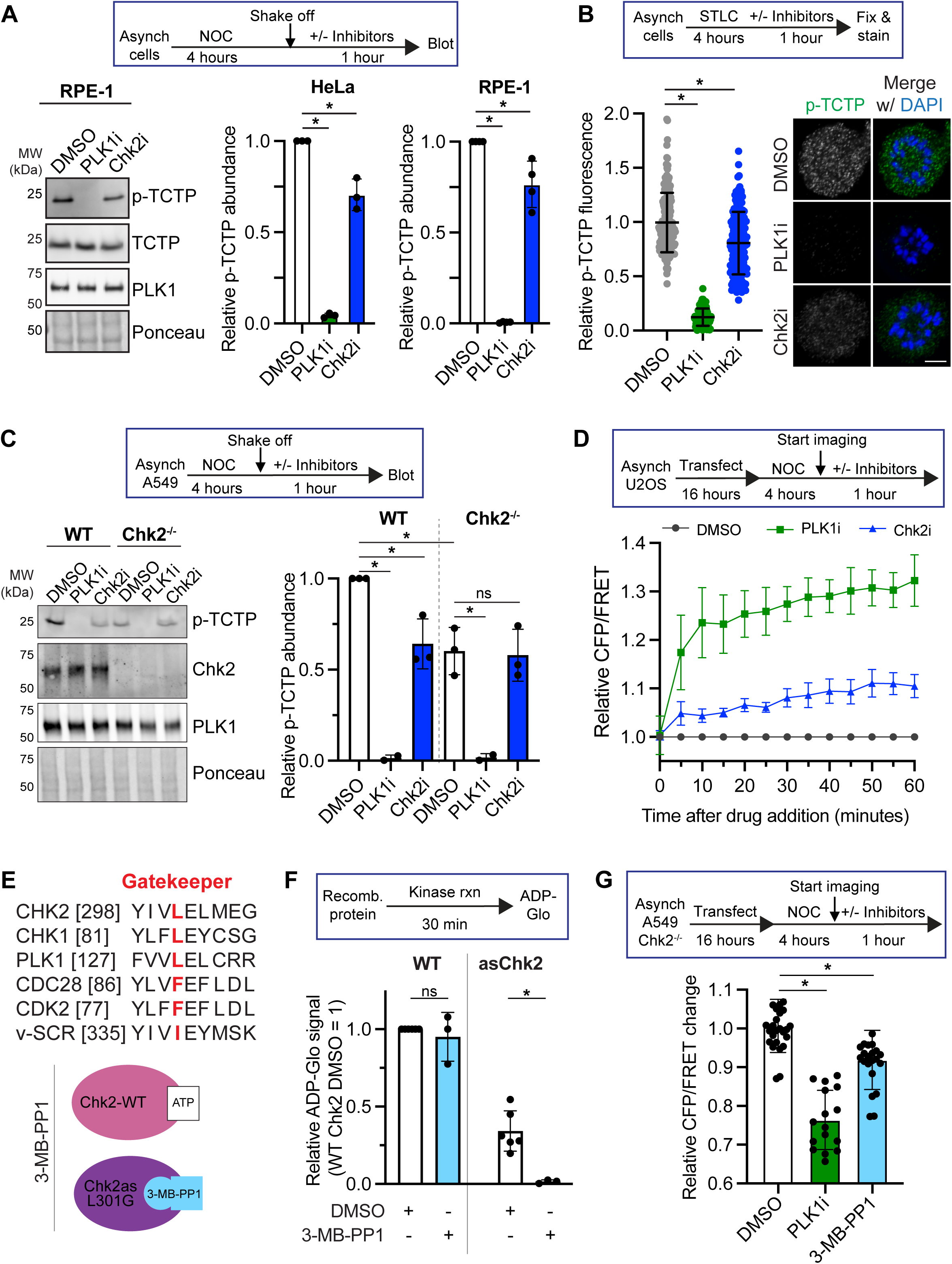
Chk2 sustains mitotic PLK1 activity. (A) Top, experimental setup. Following 4-hour treatment with 100ng/mL nocodazole (NOC), mitotic cells were collected via shake off and centrifugation. These mitotic cells were then resuspended in a smaller volume of the original media (containing NOC) and treated for 1 hour with PLK1i (1μM BI-2536), Chk2i (10μM BLM-277), or equal volume DMSO as a vehicle control. Following the inhibitor treatment, cells were collected and immunoblotted. Left, representative western blot of RPE-1 cells treated as described above. Right, quantification of three experimental replicates in cell lines as marked. Each point demarcates value from biological replicate. Error bars SD. *p<0.05, two-tailed t-test. (B) Top, experimental setup. Following 4-hour arrest in 10μM S-trityl-L-cysteine (STLC), inhibitor or DMSO was added for 1 hour. Cells were then fixed in paraformaldehyde and stained with an antibody against p-TCTP. Points are individual cells combined from 3 independent experiments, ≥115 cells per condition total. Error bars SD. *p<0.05, student’s two-tailed t-test of average normalized value from each experimental replicate. (C) A549 wild-type (WT) or Chk2^-/-^ cells were treated as in (A). *p<0.05, unpaired one-way ANOVA with Bonferroni multiple comparison correction. ns, not significant. (D) Top, experimental setup. U2OS cells were transfected with a plasmid coding for the PLK1 FRET sensor overnight and then arrested in mitosis with nocodazole. An initial measurement was made for each cell before the addition of the inhibitors. Following this timepoint, the media was replaced with fresh media containing nocodazole and PLK1i, Chk2i, or DMSO as a vehicle control. Bottom, quantification of normalized CFP/FRET values for each condition. Timepoints were taken every 5 minutes for 60 minutes. Error bars SEM of replicate averages (≥3 experimental replicates, ≥28 cells per condition). (E) Top, protein sequence alignment of indicated kinase ATP binding pocket residues. Putative gatekeeper residues are colored red. Bottom, cartoon of analog-sensitive (as) Chk2 allele. While Leucine 301 in wild-type (WT) Chk2 sterically hinders the bulky ATP-competitive inhibitor 3-MB-PP1, mutating this residue to glycine generates a larger binding pocket, rendering the enzyme sensitive to 3-MB-PP1. (F) ADP-Glo quantification of kinase activity in the presence of DMSO or 10μM 3-MB-PP1. Each point is one experimental replicate. ns, not significant, *p<0.05, student’s two-tailed t-test. Full graphs for kinase activity in the presence of varying concentrations of 3-MB-PP1 can be found in Fig. S5C. (G) Top, experimental setup as in (D). Bottom, quantification of relative change in CFP/FRET ratio 1 hour after addition of inhibitors or DMSO in cells co-expressing PLK1 FRET and asChk2-mCherry. Each point represents one cell normalized to the average value in the DMSO condition for each replicate. Error bars SD of individual cell values combined from ≥3 experimental replicates, ≥16 cells per condition total. *p<0.05, two-tailed t-test of individual cell values.

To validate the specificity of Chk2i, we tested whether Chk2i affects the activity of isolated PLK1 and Aurora A, an important regulator of PLK1 ^20, 21, 34^, *in vitro* using ADP-Glo ^50^. We did not observe significant inhibition of either kinase by Chk2i (Fig. S3A). Furthermore, we compared PLK1 activity in Chk2-proficient (wild-type, WT) and -deficient (Chk2^-/-^, Fig. S3B) A549 cells treated with a vehicle control, PLK1i, or Chk2i. Chk2^-/-^ cells have ∼30% less p-TCTP compared to WT cells, comparable to Chk2i-treated WT cells (Fig. 1C, S3B-C). Chk2i treatment did not further affect p-TCTP in Chk2^-/-^ cells, providing evidence for the specificity of the inhibitor in this context. We also observed reduced p-TCTP in nocodazole-arrested HeLa, U2OS, and A549 cells treated with a Chk2-targeting siRNA compared to a non-targeting siRNA (Fig. S3D), demonstrating that genetic approaches to disrupt Chk2 also lead to a reduction in mitotic PLK1 activity.

We validated that p-TCTP is a reliable marker of PLK1 activity with analog-sensitive (as) PLK1 RPE-1 cells in which the endogenous PLK1 locus is modified to contain mutations to the ATP binding pocket that renders the protein uniquely susceptible to inhibition with 3-MB-PP1, a bulky and nonhydrolyzable ATP analog ^6^. Using this system, we observed a complete loss of p-TCTP signal following inhibition by 3-MB-PP1, demonstrating that p-TCTP S46 is a specific output for PLK1 activity (Fig. S4A). As an additional readout of PLK1 activity, we used an engineered FRET sensor whose conformation and subsequent FRET emission is correlated with PLK1 activity ^7^ (Fig. S4B). PLK1i treatment of nocodazole-arrested U2OS and RPE-1 cells led to a ∼30% decrease in normalized CFP/FRET signal. In comparison, Chk2i treatment led to a ∼7% decrease in normalized CFP/FRET signal after one hour (Fig. 1D, S4C). These data support our observation that Chk2 inhibition leads to a partial reduction in mitotic PLK1 activity.

Finally, we sought to develop a chemical genetics approach to validate that Chk2 sustains PLK1 activity. We engineered an asChk2 allele by mutating Chk2’s gatekeeper residue, a conserved amino acid in the ATP binding pocket, from Leucine to Glycine. This mutation increases the size of the ATP-binding pocket and renders the protein uniquely susceptible to inhibition with 3-MB-PP1 (Fig. 1E). We purified WT and asChk2 (L301G) from *E. coli* (Fig. S5A-B) and determined their relative activity in the presence of DMSO or 3-MB-PP1 using ADP-Glo (Fig. S5C). Although asChk2 had moderately lower basal activity than WT Chk2, we observed that only asChk2 activity was significantly affected by treatment with 3-MB-PP1 (Fig. 1F), demonstrating the selectivity of this inhibitor for the as-kinase and our ability to use asChk2 as a genetic tool to study Chk2 function. We measured PLK1 activity in Chk2^-/-^ A549 cells that were transiently co-transfected with the PLK1 FRET sensor and mCherry-asChk2. Cells expressing asChk2 treated with 3-MB-PP1 demonstrated a significant decrease in the CFP/FRET ratio (Fig. 1G), reflecting a decrease in PLK1 activity. In contrast, cells that did not express asChk2 (mCherry-negative) had no significant change in the CFP/FRET ratio following 3-MB-PP1 treatment (Fig. S5D), indicating that this effect is specific to inhibiting asChk2. These data use a chemical genetics approach to further support our observation that Chk2 sustains PLK1 activity in mitosis.

### Chk2 activity towards PLK1 is independent of the DNA damage response pathway

We wanted to identify whether this novel function for Chk2 in regulating mitotic PLK1 activity is dependent on ATM, as it is a well-characterized Chk2 activator in response to DNA damage ^30, 51, 52^. First, we validated that U2OS and HeLa cells are ATM-proficient by treating cells with doxorubicin to induce DNA damage and ATM signaling. We observed increased phosphorylation of the canonical ATM substrate p-Chk2 S33/35 ^53^ following doxorubicin treatment, and this increase was sensitive to treatment with ATM inhibitor (ATMi, 10μM KU55933, Fig. S6A). ATM inhibition in mitotic U2OS cells did not significantly affect PLK1 activity as measured by CFP/FRET emission for the PLK1 FRET sensor or p-TCTP abundance (Fig. 2A, Fig. S6B).

**Fig. 2.**
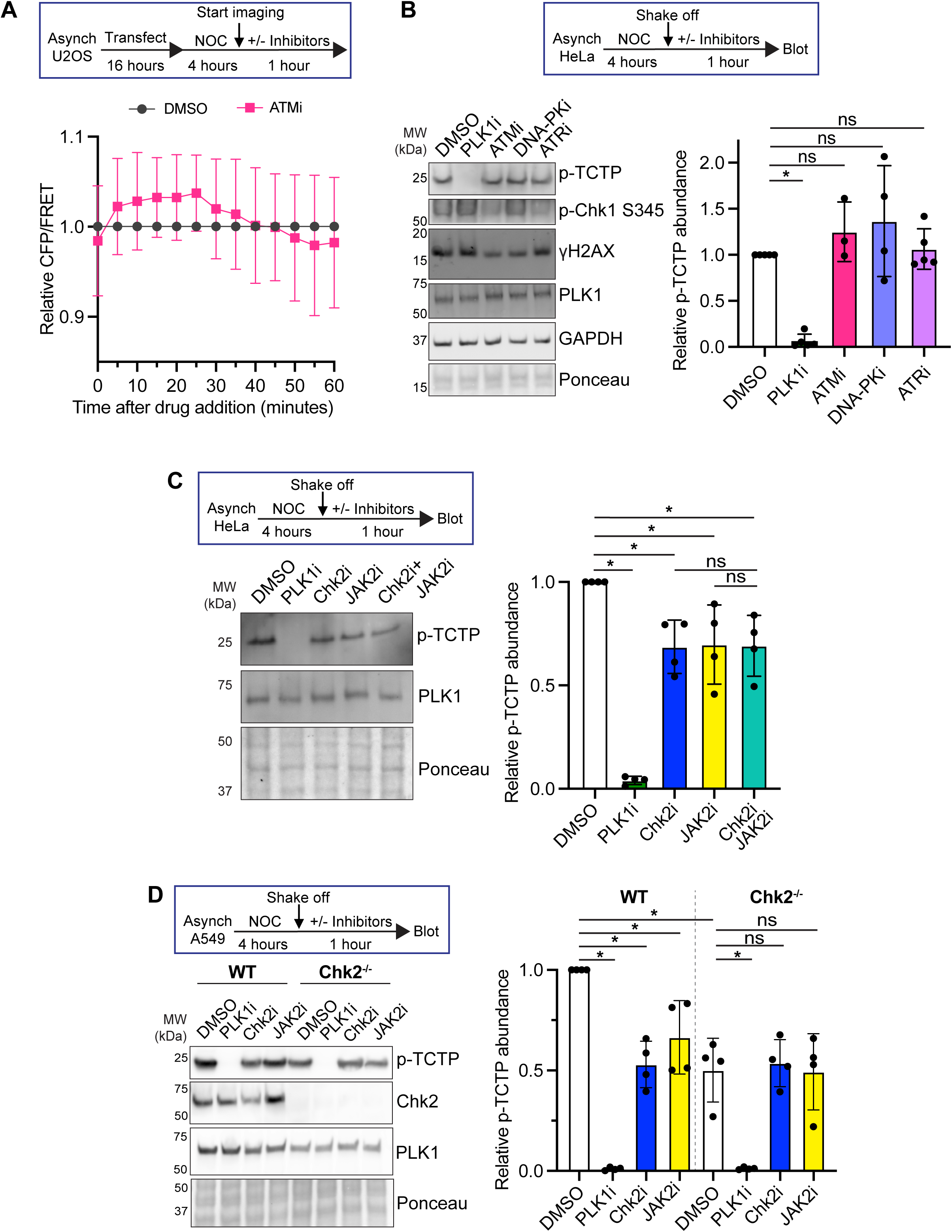
Chk2 works in concert with JAK2 to promote PLK1 activity (A) Quantification of relative change in CFP/FRET ratio 1 hour after addition of ATMi (10μM KU55933) or DMSO in nocodazole-arrested U2OS cells. Error bars SEM of replicate averages (≥3 experimental replicates, ≥56 cells total). (B) Top, experimental setup. NOC is 100ng/mL nocodazole. Following 4-hour treatment with nocodazole, mitotic cells were collected via shake off and centrifugation. These mitotic cells were then resuspended in a smaller volume of the original media (with NOC) and treated for 1 hour with PLK1i (1μM BI-2536), ATMi (10μM KU55933), DNA-PKi (5μM M3814), ATRi (10μM AZ-20), or equal volume DMSO as a vehicle control. Following the inhibitor treatment, cells were collected and immunoblotted. Right, quantification of ≥3 experimental replicates. Each point demarcates value from biological replicate. Error bars SD. *p<0.05, two-tailed t-test. (C) Top, experimental setup. Isolated mitotic HeLa cells were treated for 1 hour with PLK1i (1μM BI-2536), Chk2i (10μM BML-277), JAK2i (5μM JAK2 inhibitor IV), combination of Chk2i and JAK2i, or equal volume DMSO as a vehicle control. Following inhibitor treatment, cells were collected and immunoblotted. Left, representative western blot. Right, quantification of 4 experimental replicates. Each point demarcates the value from one experimental replicate. Error bars SD. *p<0.05, one-way anova with Bonferroni multiple comparison correction. (D) Top, experimental setup. Isolated mitotic A549 WT and Chk2^-/-^ cells were treated as in 2C. Left, representative western blot. Right, quantification of 4 experimental replicates. Each point demarcates value from biological replicate. Error bars SD. *p<0.05, one-way anova with Bonferroni multiple comparison correction.

Although ATM is the best-characterized Chk2 activator, other apical DNA damage response kinases, including ATR and DNA-PK, have also been reported to active Chk2 ^54, 55^. To test whether these kinases affect PLK1 activity, we treated isolated nocodazole-arrested HeLa cells with small molecule inhibitors of ATM, DNA-PK (DNA-PKi, 5μM M3814), and ATR (ATRi, 10μM AZ20). We did not observe any significant change in p-TCTP with these inhibitors compared to a DMSO control (Fig. 2B). In contrast, inhibitor treatment led to reduced phosphorylation of the ATM and DNA-PK substrate γH2AX and the ATR substrate phospho-Chk1 S345, demonstrating inhibitor efficacy (Fig. 2B). Together, these data demonstrate that ATM, DNA-PK, and ATR do not affect PLK1 activity in mitosis and suggest that mitotic Chk2 activity is independent of canonical upstream regulators in the DNA damage response pathway.

Interestingly, we confirmed a recent report that the nonreceptor tyrosine kinase Janus kinase 2 (JAK2) promotes mitotic Chk2 activity ^42^ and found that JAK2 inhibition (JAK2i, 5μM JAK2 inhibitor IV) also reduces PLK1 activity in nocodazole-arrested U2OS, HeLa, and A549 cells by ∼30%, comparable to the reduction we observe following treatment with Chk2i (Fig. 2C-D, Fig. S6B). Combinatorial treatment of JAK2i and Chk2i did not decrease p-TCTP beyond either inhibitor alone (Fig. 2C, Fig. S6B), suggesting that JAK2 and Chk2 work in the same pathway to promote PLK1 activity. To further validate whether JAK2’s effect on PLK1 is dependent on Chk2, we treated nocodazole-arrested A549 WT and Chk2^-/-^ cells with JAK2i and observed no additional effect on p-TCTP levels (Fig. 2D), supporting the hypothesis that noncanonical activation of Chk2 by JAK2 is important for regulating mitotic PLK1 activity.

### Chk2 directly phosphorylates PLK1 T210

To identify potential mechanisms by which Chk2 promotes PLK1 activity, we sought to determine if Chk2 modulates the known PLK1 regulatory kinases CDK1 and Aurora A. CDK1 primes PLK1 activity by phosphorylating key scaffolding proteins at the centrosome and kinetochore, which in turn recruit PLK1 through its polo-binding domains ^13, 56, 57^. One-hour treatment of mitotic U2OS cells treated with the proteasome inhibitor MG132 (20μM) and CDK1 inhibitor (CDK1i, 5μM RO-3306) led to a significant reduction in PLK1 activity, measured by p-TCTP (Fig. 3A). Combinatorial treatment with CDK1i and Chk2i in the presence of MG132 further reduced p-TCTP, suggesting that Chk2 works independently of CDK1 to promote PLK1 activity (Fig. 3A). These changes are not due to premature mitotic exit following treatment with CDK1i, as cyclin B levels were unaffected (Fig. 3A). Furthermore, Chk2i does not affect the abundance of phospho-PRC1 T481 (p-PRC1), a known mitotic CDK1 substrate ^58^ (Fig. S7A-C). These data are consistent with our model that Chk2 promotes PLK1 activity independently of CDK1.

**Fig. 3.**
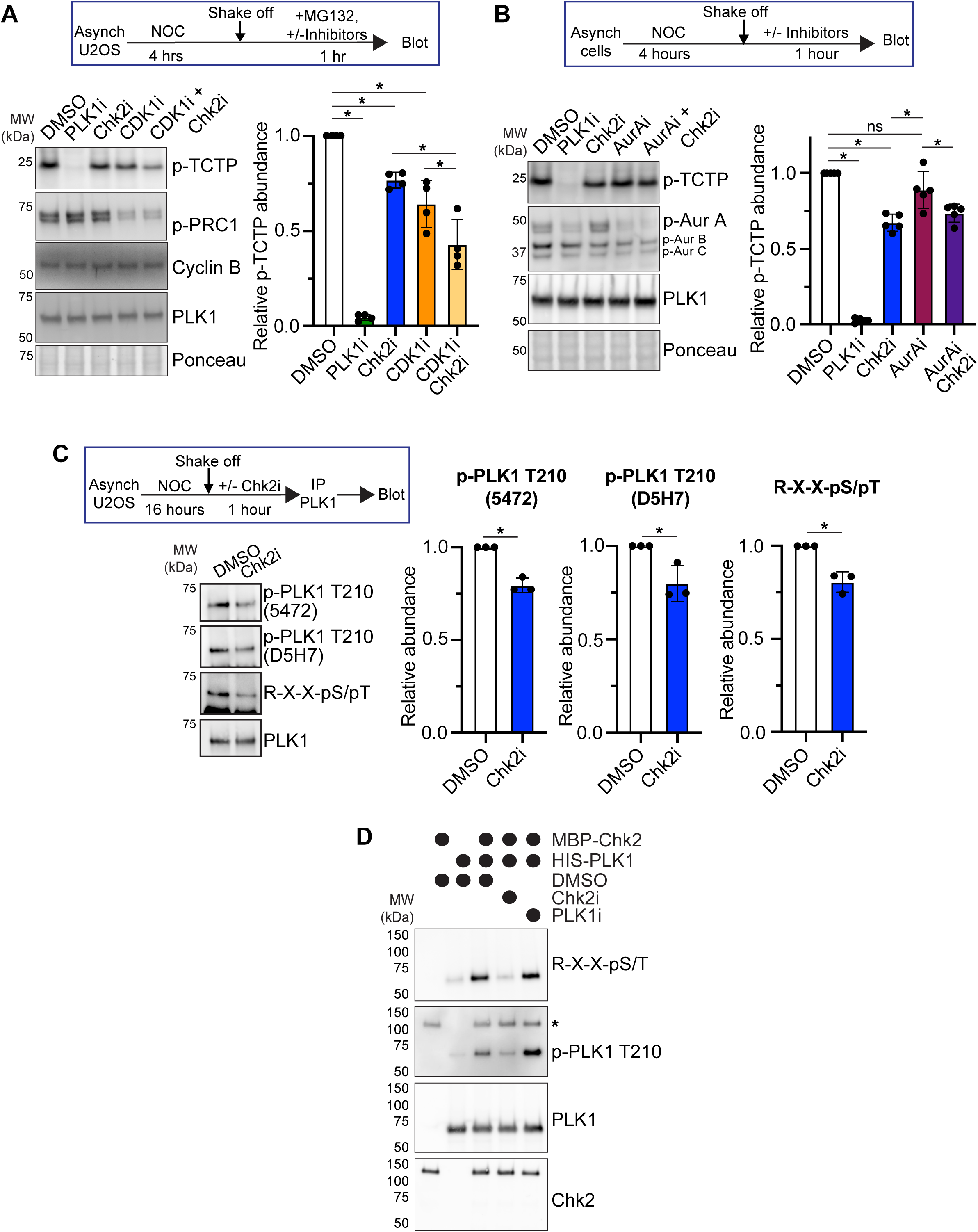
Chk2 regulates PLK1 through direct phosphorylation (A) Representative western blot (left) and quantification (right) of U2OS cells treated with MG132 (20μM) in combination with PLK1i (1μM BI2536), Chk2i (10μM BLM-277), CDK1i (5μM RO-3306), CDK1i Chk2i combination, or a DMSO vehicle control. Each point represents one experimental replicate *p<0.05, paired 1-way ANOVA with Bonferroni multiple comparisons correction. ns, not significant (p>0.05). (B) Representative western blot (left) and quantification (right) of U2OS cells treated with PLK1i (1μM BI2536), Chk2i (10μM BLM-277), AurAi (50nM MLN-8237), AurAi Chk2i combination, or a DMSO vehicle control. *p<0.05, 1-way ANOVA with Bonferroni multiple comparisons correction. ns, not significant (p>0.05). (C) Experimental setup (top) and representative western blot (left) of PLK1 immunoprecipitated from mitotic HeLa cells treated with DMSO or Chk2i. Right, quantification of phospho-antibody signal normalized to total PLK1 abundance. Each point represents the value from one experimental replicate. *p<0.05, two-tailed t-test (D) *In vitro* kinase assay with recombinant HIS-PLK1, MBP-Chk2 incubated with DMSO, PLK1i, or Chk2i as marked. Recombinant proteins were incubated in kinase assay buffer in a 1:1 molar ratio with 500μM ATP for 30 minutes at 30°C with gentle agitation. The reaction was quenched with 5mM EDTA before being combined with denaturing sample buffer and analyzed via western blotting. p-PLK1 T210 antibody shown is Cell Signaling Technology D5H7. *, nonspecific recognition of MBP-Chk2

Aurora A, alongside its cofactor Bora, activates PLK1 in G2 by phosphorylating PLK1 T210, a residue in PLK1’s T-loop essential for robust PLK1 activity ^20–22^. Bora is mostly degraded at mitotic entry, and Aurora A’s contribution to sustaining PLK1 activity in mitosis is limited to ∼15-20% ^33, 34^. Consistent with prior reports ^34^, acute Aurora A inhibition (AurAi, 50nM MLN-8237) of mitotic U2OS cells led to a ∼15% loss of PLK1 activity (Fig. 3B). Co-treatment of mitotic cells with AurAi and Chk2i did not further decrease p-TCTP beyond Chk2i, indicating that Aurora A activity may contribute to some of the reduction in PLK1 activity following treatment with Chk2i. However, the total decrease in p-TCTP with Chk2i was greater than AurAi alone, indicating that changes in Aurora A activity are not sufficient to fully explain the change in PLK1 activity we observe with Chk2i. In support of this model, we did not observe significant changes in the abundance of phospho-Aurora A T288 (p-AurA), a marker of active Aurora A, when we blotted an antibody that recognizes phosphorylated Aurora A/B/C kinases, suggesting that Chk2 is not exclusively working though Aurora A to regulate PLK1 (Fig. 3B, Fig. S7). Taken together, these data demonstrate that Chk2 does not sustain mitotic PLK1 activity exclusively by modulating the known PLK1 regulators CDK1 and Aurora A.

Because PLK1 activity is regulated through phosphorylation ^21, 22, 34, 59–62^, we tested the hypothesis that PLK1 phosphorylation is affected following acute treatment with Chk2i. We detected phosphorylated PLK1 with a PhosTag gel, where the migration of phosphorylated proteins is slowed, resulting in apparently higher molecular weight species that are not present in a SDS-PAGE gel ^63^ (Fig. 3C). We observed a ∼25% reduction in the proportion of phosphorylated PLK1 in mitotic HeLa cells treated with Chk2i compared to DMSO (Fig. S8A), suggesting that Chk2 promotes PLK1 phosphorylation in mitosis.

One of the most important and well-studied regulatory phosphorylation sites on PLK1 is T210, whose phosphorylation is both necessary and sufficient to promote PLK1 activity ^21, 22, 34, 59^. We tested whether PLK1 T210 phosphorylation is reduced in mitotic cells following Chk2i treatment by immunoprecipitating PLK1 from mitotic U2OS cells and probing with antibodies that specifically recognize PLK1 T210 phosphorylation (p-T210, Fig. S8B). Treatment with Chk2i significantly (∼25%) reduced p-T210 normalized to the total immunoprecipitated PLK1 (Fig. 3C), and this effect was reproducible with two independent antibodies (Cell Signaling Technology #5472 and #D5H7). Likewise, we probed the immunoprecipitated protein with an antibody that recognizes phosphorylated Serine or Threonine residues with an Arginine at the -3 position (R-X-X-pS/pT), as PLK1 T210 matches this motif (R-K-K-<u>T</u>). We observed that Chk2-treated cells had a similar decrease in signal from the phospho-antibody, consistent with the observation that this phosphorylation site is sensitive to treatment with Chk2i.

Interestingly, Chk2’s substrate consensus motif is highly basophilic ^64, 65^, and PLK1 T210 scores above the 97^th^ percentile as a computationally predicted Chk2 substrate (that is, PLK1 T210 scores more favorably as a Chk2 substrate than 97% of putative phosphorylation sites in the phosphoproteome) ^64^. This prompted our hypothesis that Chk2 may directly phosphorylate PLK1 T210 to sustain its activity in mitosis. We sought to determine whether Chk2 can directly phosphorylate PLK1 by performing an *in vitro* kinase assay where we co-incubated catalytically active MBP-Chk2 and HIS-PLK1. When we probed the resulting kinase assay with R-X-X-pS/pT and p-T210 antibodies, we observed increased signal at PLK1’s molecular weight compared to PLK1 alone (Fig. 3D). This signal was sensitive to treatment with Chk2i, but not PLK1i, suggesting that Chk2 phosphorylates PLK1. We performed a similar kinase reaction with active or kinase-dead (KD) Chk2 in which T383 and T387 are mutated to Alanine to abrogate activity^66^. Similarly, only the active form of Chk2 was able to phosphorylate PLK1 (Fig. S8C). Finally, we used Parallel Reaction Monitoring (PRM)-based mass spectrometry to measure the abundance of phospho-PLK1 T210 following co-incubation of active GST-Chk2 and HIS-PLK1 in the presence of DMSO or Chk2i, as in Fig. 3D. Using a synthetic heavy-labelled standard peptide, we verified the existence of phospho-PLK1 T210 in the kinase reaction and calculated that this phospho-peptide was 1.8 times more abundant in the DMSO-treated reaction than in the reaction with Chk2i (Fig. S8D). Together, these data support a model in which Chk2 directly phosphorylates PLK1 T210 to promote PLK1 activity in mitosis.

### PLK1 and Chk2 activity promote proper mitotic progression

PLK1 promotes proper mitotic progression through its roles in centrosome maturation, bipolar spindle formation, kinetochore-microtubule attachment, chromosome segregation, and cytokinesis ^5–12^. We hypothesized that the partial reduction in PLK1 activity following Chk2i may induce defects in some of these processes, as previous work has identified that these PLK1 functions are differentially sensitive to partial inhibition ^8^. To tease apart the PLK1-dependent and -independent effects of mitotic Chk2 inhibition, we identified a dose of PLK1i (5nM) that simulates the ∼30% reduction in p-TCTP that we observe following Chk2i in nocodazole-arrested RPE-1 cells (Fig. S9A). 5nM PLK1i also caused a similar decrease in nocodazole-arrested U2OS cells (Fig. S9B) and in asynchronous prometaphase and metaphase RPE-1 cells (Fig. 4A-B).

**Fig. 4.**
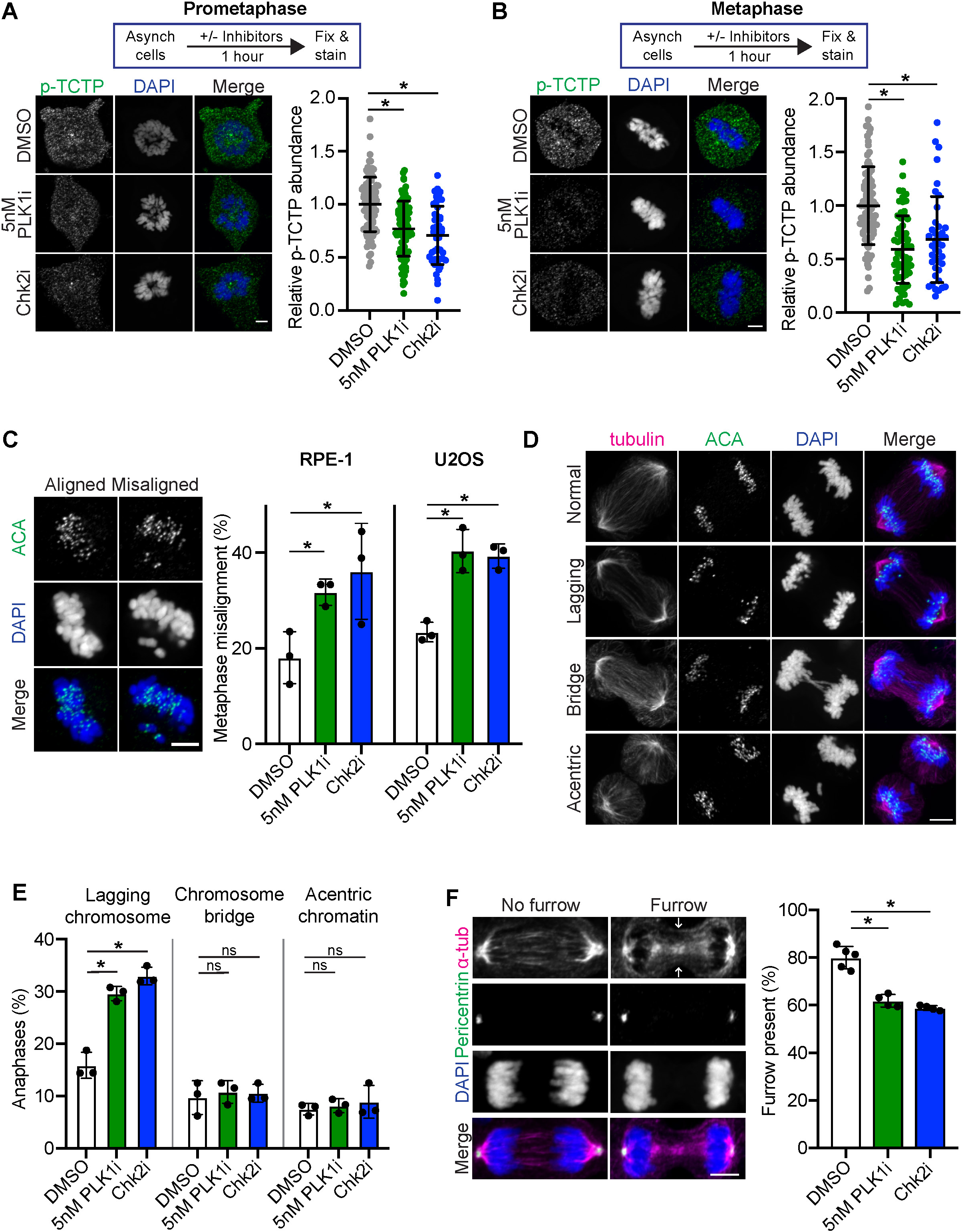
Chk2 inhibition and partial PLK1 inhibition induce mitotic defects (A-B) Left, representative image of prometaphase (A) or metaphase (B) RPE-1 cells stained for phospho-TCTP S46 (p-TCTP) following 1-hour treatment with 5nM BI-2536 (PLK1i), Chk2i (10μM BLM-277), or a DMSO vehicle control. Images are single z-planes for each channel. Scale bar, 5 μm. Right, quantification of prometaphase (A) or metaphase (B) p-TCTP signal taken from single z-slice. Each point represents one cell normalized to the average value in the DMSO condition for each replicate. Error bars SD of individual cell values combined from 3 experimental replicates, ≥48 cells (prometaphase) or ≥40 cells (metaphase) total. *p<0.05, two-tailed t-test of replicate mean values. (C) Left, representative image of U2OS cells with aligned or misaligned chromosomes. ACA, anti-centromere antibody. Scale bar, 5 μm. Right, quantification of misalignment rate in RPE-1 (left) or U2OS (right) cells following 1-hour treatment with inhibitors or DMSO control. Each point represents the rate from an experimental replicate (U2OS cells >50 cells per replicate per condition, >280 cells total per condition). *p<0.05, two-tailed t-test of replicate mean values. (D) Representative z-projected images of U2OS cells with mitotic defects. Scale bar, 5μm. (E) Quantification of mitotic defects in U2OS cells treated for 1 hour with inhibitors or vehicle control. Each point represents the rate from an experimental replicate (≥180 anaphases per condition total). *p<0.05, two-tailed t-test of replicate averages. ns, not significant (p>0.05). (F) Left, representative image of telophase RPE-1 cells stained for alpha tubulin and pericentrin following 1-hour treatment inhibitors or a DMSO vehicle control. Images are maximum intensity z-projections for each channel. Scale bar, 5μm. Right, quantification of the rate of cytokinetic furrow formation. Each point represents the rate from an experimental replicate (≥97 cells per replicate). *p<0.05, two-tailed t-test of replicate averages.

PLK1 localizes to, and is active at, the centrosome and kinetochore, and its localization is partially dependent on its kinase activity ^14, 67^. We treated asynchronous RPE-1 cells with 5nM PLK1i, Chk2i, or DMSO for 1 hour and measured PLK1 localization to centrosomes, marked by pericentrin, and kinetochores, marked by anti-centromere antibody (ACA). Importantly, this experimental setup with acute, 1-hour inhibitor treatment provides high temporal resolution to differentiate the mitotic and interphase effects of perturbing kinase signaling. Cells treated with 5nM PLK1i or Chk2i had reduced PLK1 localization to kinetochores and centrosomes in both prometaphase and metaphase (Fig. S10A-D). These data suggest that Chk2 activity is crucial to maintaining proper PLK1 activity and localization at both the kinetochore and centrosome, and that this effect is not prophase-specific.

We sought to identify the effects of acute Chk2 inhibition and partial PLK1 inhibition on proper mitotic progression. PLK1 promotes proper chromosome alignment in metaphase by phosphorylating proteins at the centromere and kinetochore that regulate the attachment to spindle microtubules and protect chromatin against microtubule pulling forces ^7, 11, 16^. We observed that RPE-1 and U2OS cells treated for 1 hour with Chk2i or 5nM PLK1i had significant defects in metaphase chromosome alignment (Fig. 4C), a prerequisite for accurate chromosome in anaphase ^68^. We observed a similar increase in metaphase misalignment in asynchronous Chk2^-/-^, compared to WT, A549 cells (Fig. S11A). Consistent with our model in which JAK2 acts as an upstream regulator of Chk2 in mitosis, treatment of WT, but not Chk2^-/-^, A549 cells with JAK2i also led to an increase in metaphase chromosome misalignment compared to DMSO (Fig. S11A).

An increased rate of anaphase chromosome missegregation is a hallmark of many kinds of cancers and is correlated with poor patient prognosis ^1–4^. We tested whether partial reduction in PLK1 activity through Chk2i or 5nM PLK1i would affect chromosome segregation fidelity. RPE-1 and U2OS cells acutely treated with Chk2i or 5nM PLK1i had increased rates of whole-chromosome missegregation, or lagging chromosomes, compared to vehicle-treated cells (Fig. 4E, S11A). In contrast, inhibitor treatment did not increase the rates of acentric chromatin or chromatin bridges, which are caused by unrepaired DNA double-stranded breaks or under-replicated DNA, respectively (Fig. 4E, S11A). We also observed that Chk2^-/-^ cells had significantly more lagging chromosomes compared to WT cells (Fig. S11C). Acute treatment with JAK2i phenocopied this effect in WT cells and did not further increase the rate of lagging chromosomes in Chk2^-/-^ cells (Fig. S11C). Consistent with our immunofluorescence data, live-cell imaging of H2B-GFP-expressing HeLa cells treated in prophase with 5nM PLK1i, Chk2i, or JAK2i caused a ∼2-3-fold increase in the rate of chromosome missegregation compared to DMSO-treated cells (Fig. S11D-E). Together, these data demonstrate that mitotic JAK2 and Chk2 activity are essential to promote proper chromosome segregation and protect genome stability through PLK1.

PLK1 activity is also important for cytokinesis at the conclusion of mitosis ^6, 8^, and failure to properly complete cytokinesis can result in tetraploidy. We sought to determine if cytokinesis is also sensitive to Chk2 and partial PLK1 inhibition. Asynchronous RPE-1 cells treated with either Chk2i or 5mM PLK1i exhibited a decreased percentage of telophase cells with a cytokinetic furrow, measured by constriction of α-tubulin between the dividing chromatin (Fig. 3F). This effect is not due to defects in spindle elongation, as the average distance between centrosomes was unchanged in Chk2i- and 5mM PLK1i-treated cells (Fig. S11F-G). These data suggest that Chk2 activity is crucial for proper PLK1 function in distinct contexts, and that even modest reductions in PLK1 activity may be deleterious to genome stability and mitotic progression. This work also highlights that some of the mitotic defects previously observed following Chk2 perturbation ^37–45^ may be due to a bona fide mitotic function rather than the result of unresolved DNA damage in interphase.

### Chk2 deficiency sensitizes cells to treatment with low-dose PLK1 inhibitors

PLK1 activity is essential for timely mitotic division, making it an attractive therapeutic target in cancer therapies. However, PLK1 inhibitors have thus far been unsuccessful as a clinical monotherapy, in part because of on-target toxicity ^69^. We hypothesized that Chk2 deficiency, which we have shown reduces mitotic PLK1 activity, could increase cellular sensitivity to low-dose PLK1 inhibitors. We treated WT and Chk2^-/-^ A549 cells for 3 days with a titration of PLK1i and measured cell viability with crystal violet. Chk2^-/-^ cells were more sensitive to PLK1i compared to WT cells at several concentrations, most prominently around 5-10nM (Fig. 5A). Consistent with these data, knocking down Chk2 with a siRNA (Fig. 5B) significantly increased sensitivity to 2.27nM PLK1i compared to a non-targeting siRNA control (Fig. 5C, S12A-C). This observation is further supported by a recent study where Chk2 knockout was identified in an unbiased genome-wide screen to confer vulnerability to low-dose PLK1 inhibition ^18^.

**Fig. 5.**
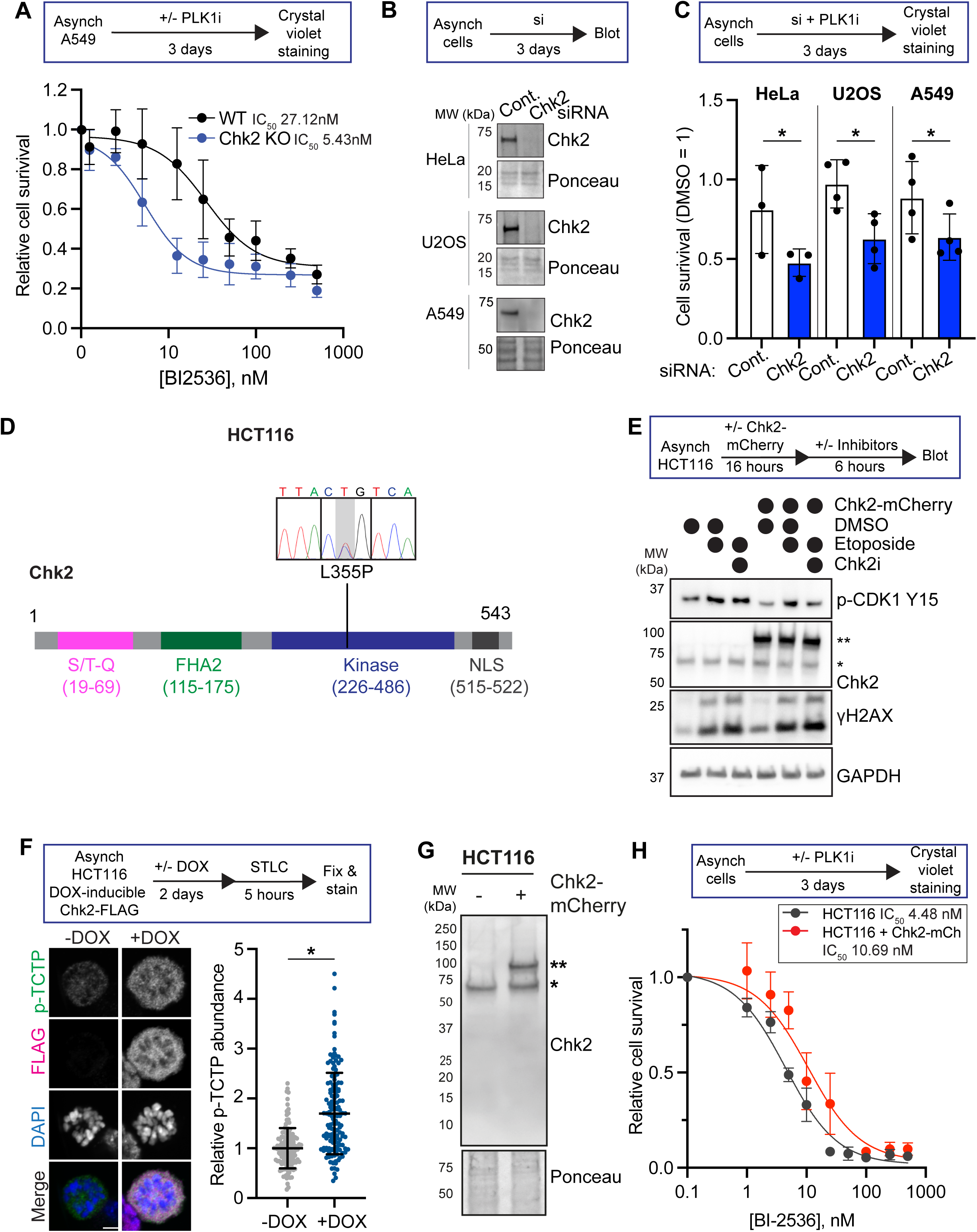
Chk2 deficiency sensitizes cells to PLK1 inhibition (A) Top, experimental setup. Bottom, relative cell viability (DMSO-treated cells = 1) following 3-day treatment with PLK1 inhibitor BI-2536. Points represent average of ≥3 experimental replicate values averaged from technical triplicates. Error bars SEM of replicate averages. (B) Representative western blots of cells treated for 3 days with either a nontargeting siRNA (control) or siRNA against Chk2 in Hela, U2OS, and A549 cells. (C) Quantification of cell viability measured by crystal violet staining after 3 days of co-incubation with siRNA (either nontargeting control or against Chk2) and 2.27nM PLK1 inhibitor BI-2536. Full curves for BI-2536 sensitivity can be found in Fig. S12A-C. Each point marks the average of technical triplicates within one experimental replicate normalized to signal from DMSO-treated cells. Error bars SD of replicate averages. *p<0.05, paired two-tailed t-test of replicate mean values. (D) Map of Chk2 protein with indicated functional domains including S/T-Q cluster, phospho-peptide-binding and dimerization interface FHA2 domain, kinase domain, and nuclear localization sequence (NLS). Top, trace files of associated genomic loci amplified from HCT-116 cells. Gray shading indicates the heterozygous L355P mutation. (E) Top, experimental setup. Bottom, representative western blot of HCT-116 cells with or without the Chk2-mCherry rescue construct challenged with the DNA damaging agent etoposide (Etop, 10μM) or in combination with Chk2i (10μM BLM-277) for 6 hours. (F) Left, quantification of p-TCTP abundance in STLC-arrested HCT-116 cells transduced with a doxycycline (DOX)-inducible Chk2-3X FLAG construct. Cells were either treated with 0.1μM DOX or untreated for 48hrs prior to STLC addition. Each point represents one cell normalized to the average value in the untreated condition for each replicate. Error bars SD of individual cell values combined from 3 experimental replicates, ≥131 cells total. *p<0.05, two-tailed t-test of replicate mean values. Right, representative immunofluorescence image from single z-plane. Scale bar, 5μm. Relative expression of Chk2 construct following induction can be found in Fig. S12D. (G) Left, representative western blot of HCT-116 cells with or without expression of the Chk2-mCherry rescue construct. Right, quantification of relative cell viability in HCT-116 cells following 3-day treatment with BI-2536 titration. Points represent average of ≥3 experimental replicate values averaged from technical triplicates. Error bars SEM of replicate averages.

Finally, we wanted to determine if we could rescue PLK1 inhibitor sensitivity in cells carrying an endogenous Chk2 mutation. HCT-116 cells have a heterozygous L355P mutation in Chk2’s kinase domain that is predicted to damage Chk2 kinase activity ^70^ (Fig. 5D). We tested the functional status of Chk2 in this HCT-116 parental cell line and cells expressing an mCherry-tagged rescue construct by inducing DNA damage with the topoisomerase II inhibitor etoposide and measuring CDK1 Y15 phosphorylation (p-Y15), a canonical output of Chk2 activity ^70^. While parental HCT-116 cells had increased p-Y15 signal following DNA damage, likely because of compensation or redundancy with the DNA damage response kinase Chk1 ^28, 71^, they were insensitive to treatment with Chk2i (Fig. 5E). In contrast, p-Y15 abundance was sensitive to treatment with Chk2i in the rescue cell line, confirming the functional status of the Chk2 rescue line.

We measured PLK1 activity in HCT-116 cells in cells transduced with a doxycycline (dox)-inducible Chk2-FLAG construct. Following 2 days of dox induction, cells expressing Chk2 had significantly higher p-TCTP levels (Fig. 5F, S12D), demonstrating that Chk2 functional status modulates PLK1 activity, and that this mitotic Chk2-PLK1 pathway may contribute to PLK1 inhibitor sensitivity. When we treated parental HCT-116 and Chk2 rescue cells with PLK1i and measured viability, we observed that Chk2-expressing cells were less sensitive to low-dose PLK1 inhibition (Fig. 5G). Together, these data demonstrate that Chk2 status may be a predictive biomarker of PLK1i sensitivity, and that Chk2 deficiency could be a therapeutic niche in which PLK1 inhibitors are an effective cancer therapeutic strategy.

## Discussion

PLK1 is a well-established safeguard of the genome in part through its multiple mitotic functions. Tight control of PLK1’s spatiotemporal localization, as well as its precise activity, is essential for proper chromosome segregation. Chk2 protects genome stability through its role in the DNA damage response pathway, where it counteracts CDK1 and PLK1 activity as part of the G2/M checkpoint. Our data reveal an unexpected function for Chk2 to sustain PLK1 activity in mitosis. Disruptions to this pathway lead to mitotic errors, including impaired chromosome alignment, chromosome segregation, and cytokinesis, which are linked to cancer initiation and progression ^1–4, 72, 73^. Our work provides important context for understanding how Chk2 mutations, which are penetrant risk alleles for several types of cancer ^74–76^, influences disease initiation and progression.

This study highlights how well-characterized interphase signaling pathways, including Chk2 activation and G2/M cell cycle arrest by the DNA damage response pathway, are rewired in mitosis. While Chk2 activation in interphase strictly relies on activation by apical DNA damage response kinases ^30, 51, 54, 55^, we identified that these proteins are dispensable for the Chk2-PLK1 pathway described here. Instead, we describe how the nonreceptor tyrosine kinase JAK2, which canonically functions in innate immune signaling, is important for Chk2-mediated PLK1 activity. Our work builds on that of Chowdhury et al ^42^ to show, to our knowledge, the first evidence of Chk2 activation and function independent of other members of the DNA damage response pathway.

The DNA damage response pathway antagonizes mitotic kinase activity in interphase by inactivating CDK1, Aurora A, and PLK1, but our work, along with the work of others ^40, 44, 77, 78^, has demonstrated that this relationship is inverted in mitosis, where members of the DNA damage response pathway support mitotic kinase signaling. The molecular logic for this flipped relationship is incompletely understood but may be due in part to mitotic degradation of key players, including Bora and the CDK1-inhibiting kinase Wee1 ^25, 33^. Another possibility by which mitosis-specific signaling interactions may occur is that the spatial segregation of cytosolic and nuclear components in interphase is absent in mitosis. Nuclear envelope breakdown in early mitosis may allow for nuclear proteins, such as Chk2, to interact with cytoplasmic proteins in a phase-specific manner, including JAK2 and PLK1. PLK1 is mostly cytosolic in interphase and translocates to the nucleus only when active ^79^, thereby restricting the interaction between Chk2 and inactive cytosolic PLK1. After nuclear envelope breakdown, however, Chk2 is no longer spatially segregated from inactive PLK1 and may directly phosphorylate it to promote its activity.

We hypothesize that the rewired relationship between Chk2 and PLK1 in mitosis may trigger a switch between different modes of PLK1 regulation. The G2 activation of PLK1 by Aurora A-Bora is a highly sensitized “bistable switch” whereby both kinases promote the others’ activity, likely as a mechanism to ensure high activity and commitment to enter mitosis ^26^. This bistability, however, is lost in mitosis. Both by inhibiting Chk2 and through partial inhibition of PLK1 with low-dose PLK1 inhibitors, we observe that cells exhibit a stable and partial reduction in PLK1 activity. As PLK1 has been shown to negatively regulate Chk2 activity ^80^, this could be due to the switch from a positive feedback loop in G2 to a negative feedback relationship in mitosis. This negative regulation may be particularly important in contexts where PLK1 activity must be precisely fine-tuned, such as controlling kinetochore-microtubule attachments ^7^, where both hyper- and hypo-activity negatively impact proper mitotic progression ^81–84^. Future work will be needed to identify the specific populations of PLK1 regulated by Chk2. Together, our work shows that regulation of PLK1 in mitosis is more complex than previously appreciated and that a better understanding of the pathways that fine-tune PLK1 activity may identify novel therapeutic vulnerabilities for cancer treatment.

## Acknowledgements

We thank Dr. Mark Burkard for generously providing RPE-1 asPLK1 cells, Dr. Jesse Reinhart for providing the wild-type and kinase-dead Chk2 protein, Dr. Claudio Alarcon for pGEX6p-1 plasmid, and Drs. Michael Yaffe, Kevin Janes, Michael Lampson, and Bob Weinberg for reagents.

## Author contributions

E.B. and L.K. designed the study. E.B., I.T., C.R., Y.K.J., J.P., W.L., and Y.L., performed the experiments and analyses. E.B and L.K. prepared the manuscript with contributions from all authors.

## Data and materials availability

All data and unique reagents generated for this manuscript are available upon reasonable request.

## Supplemental Figure Legends

**Fig. S1.**
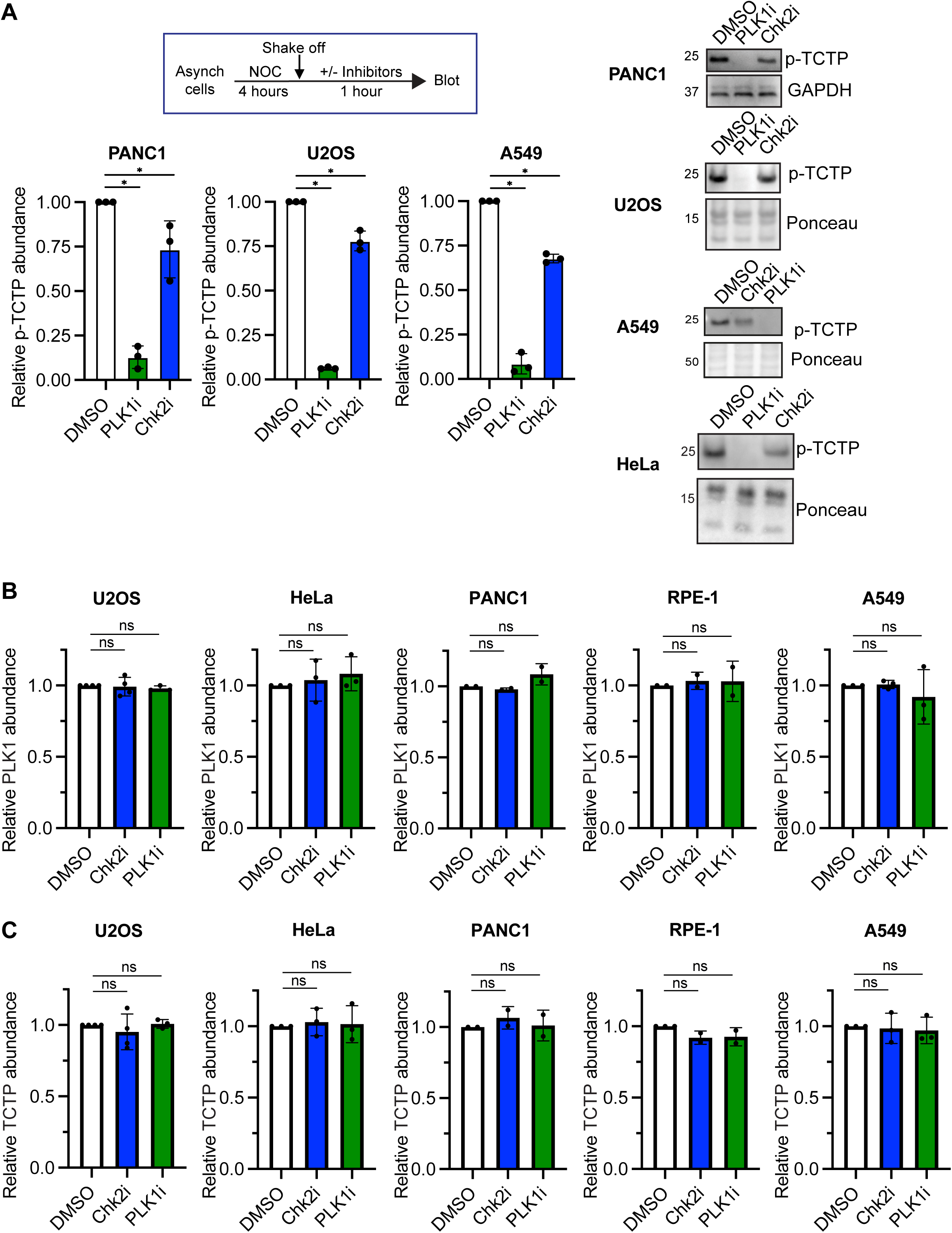
(A) Top, experimental setup. Following 4-hour treatment with 100ng/mL nocodazole (NOC), mitotic cells were collected via shake off and centrifugation. These mitotic cells were then resuspended in a smaller volume of the original media (containing NOC) and treated for 1 hour with PLK1i (1μM BI-2536), Chk2i (10μM BLM-277), or equal volume DMSO as a vehicle control. Following the inhibitor treatment, cells were collected and immunoblotted. Bottom, quantification of three experimental replicates in cell lines as marked. Each point demarcates value from biological replicate. Error bars SD. *p<0.05, two-tailed t-test. Right, representative western blots of PANC-1, U2OS, A549, and HeLa cells treated as above. The ponceau image for HeLa cells is from the same experiments as in Fig. S7A. The p-TCTP and ponceau images for U2OS cells are the same as experiment in Fig. S6B. (B-C) Quantification of western blot signal of PLK1 (B) and TCTP (C) from cells treated as in A. Each point represents value from one experimental replicate. ns, not significant (p>0.05) two-tailed t-test

**Fig. S2.**
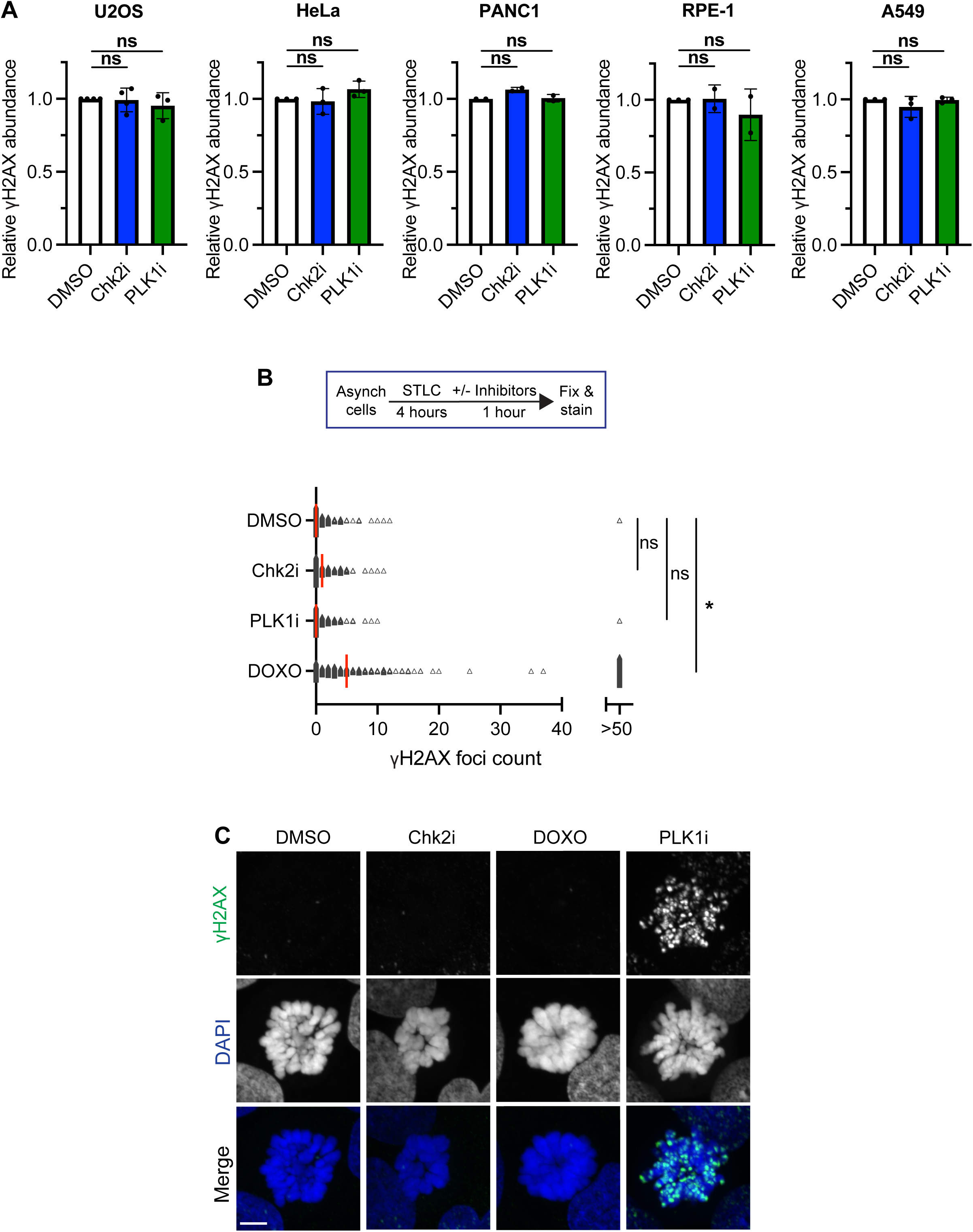
(A) As in Fig. S1B-C, quantification of γH2AX signal on western blot from cells treated as in Fig. 1A. Each point represents value from one experimental replicate. ns, not significant (all cell lines and conditions) (B) Cumulative γH2AX foci counts of STLC-arrested U2OS cells treated with DMSO, Chk2i, PLK1i, or doxorubicin (DOXO, 2μM). Each point represents one cell. Data are from 3 experimental replicates, ≥243 cells per condition total. The red line denotes the median of the data. *p<0.05; ns, not significant, two-tailed t-test of all points. (C) Representative maximum intensity projection immunofluorescence images of U2OS cells treated as in (B). Scale bar, 5μM.

**Fig. S3.**
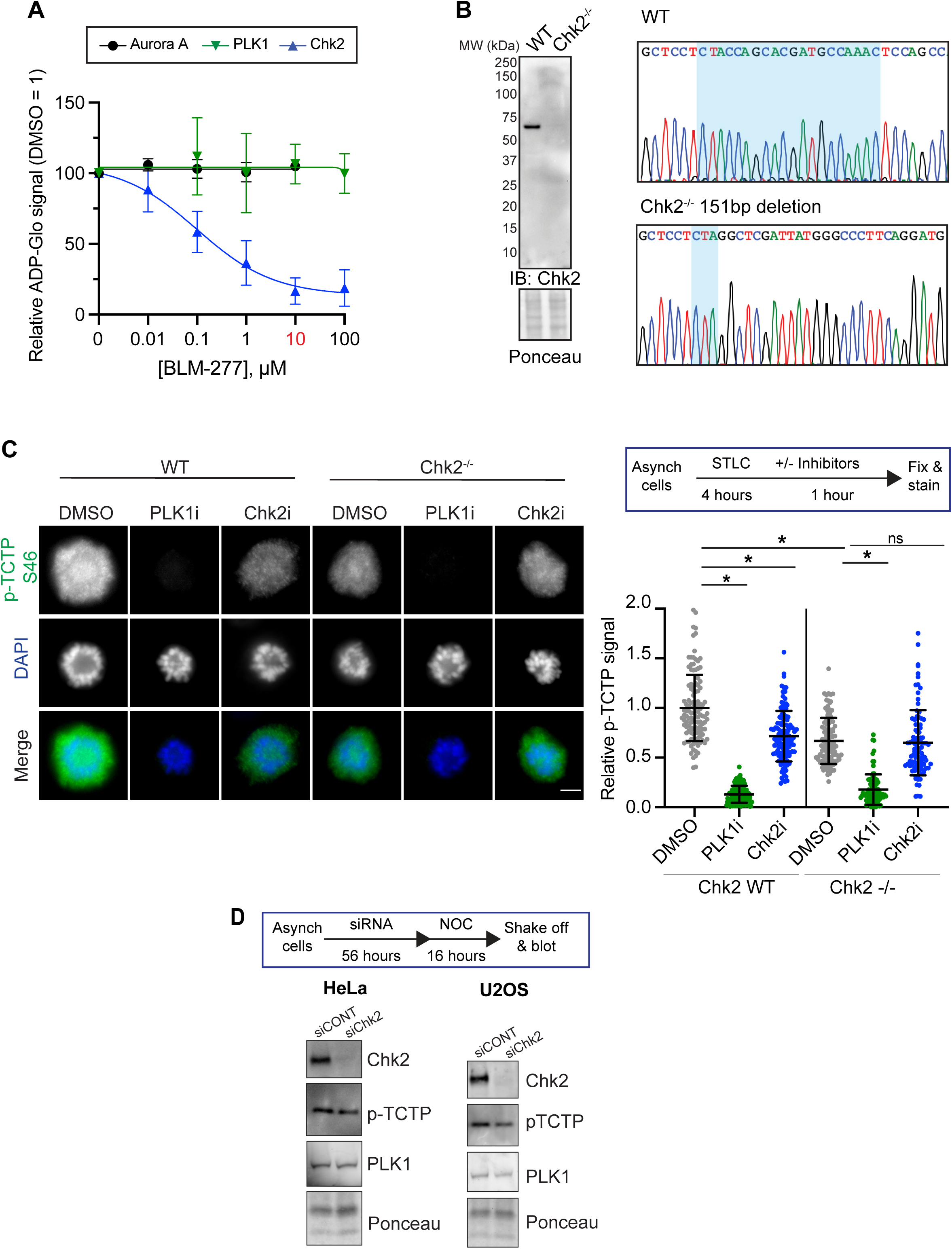
(A) ADP-Glo quantification of kinase activity in the presence of DMSO or varying concentrations of Chk2i BLM-277. Error bars SEM of ≥3 experimental replicates. (B) Left, western blot of cell extract from wild-type (WT) or Chk2^-/-^ A549 cell lines. Right, trace files of *Chek2* gene in WT or Chk2^-/-^ cells, validating the presence of a homozygous frameshift mutation in mutant cells. Light blue box is sgRNA sequence. (C) Left, representative single z-plane images of A549 wild-type (WT) or Chk2^-/-^ cells treated with inhibitors as in Fig. 1B. Scale bar, 5μm. Right, quantification of p-TCTP fluorescence intensity. Each point denotes the value from one cell normalized to the mean DMSO value for that replicate. Points are combined across 3 independent replicates. *p<0.05, 1-way ANOVA with Bonferroni correction of mean value from each experimental replicate. ns, not significant (p>0.05) (D) Top, experimental setup. Asynchronous cells were treated for 3 days with either nontargeting control or Chk2-targeting siRNA. 16 hours before the 3-day treatment was finished, cells were arrested in nocodazole (NOC). Mitotic cells were collected via shake off and immunoblotted.

**Fig. S4.**
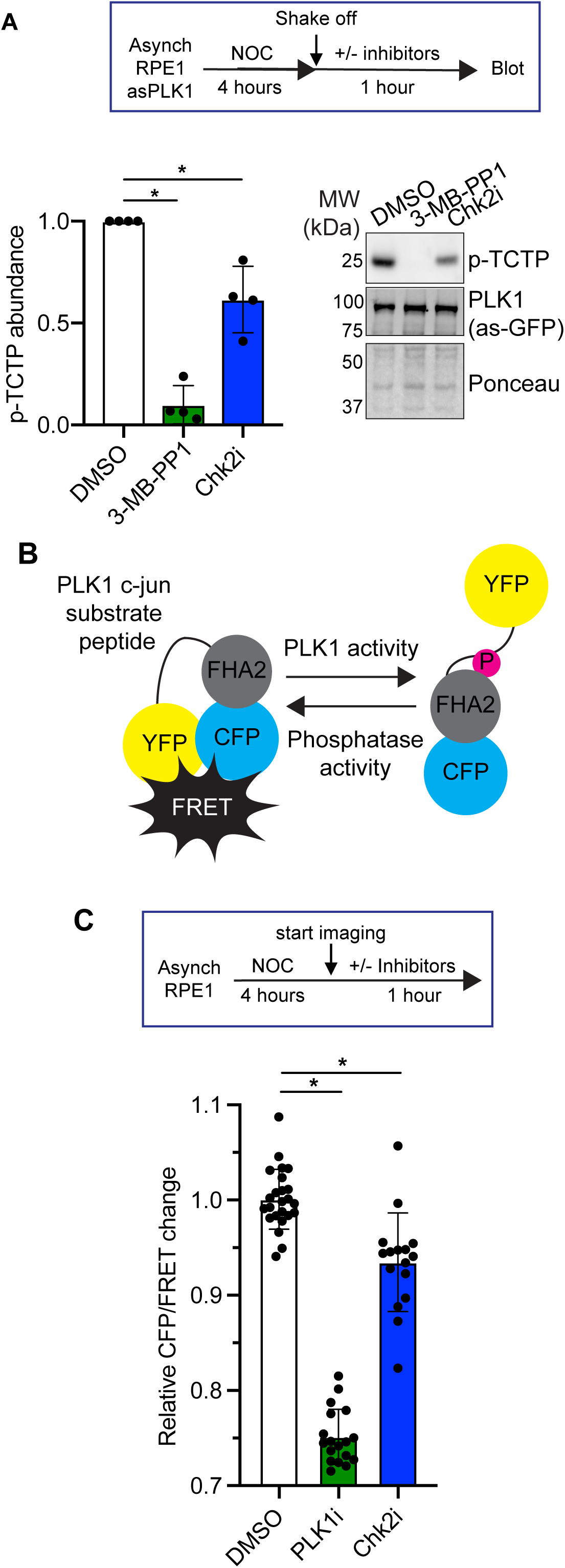
(A) Top, experimental design. Left, quantification of p-TCTP abundance in asPLK1-GFP RPE-1 cells treated with 10μM 3-MB-PP1 or Chk2i (10μM BML-277) for 1 hour. Each point represents one experimental replicate, *p<0.05, unpaired two-tailed t-test. Right, representative western blot. (B) Diagram of PLK1 FRET sensor. Increased FRET is correlated with reduced PLK1 activity. (C) Top, experimental setup as in Fig. 1D. Bottom, quantification of relative change in CFP/FRET ratio 1 hour after addition of inhibitors or DMSO. Each point represents one cell normalized to the average value in the DMSO condition for each replicate. Error bars SD of individual cell values combined from 3 experimental replicates, ≥16 cells per condition. *p<0.05, two-tailed t-test of individual cell values.

**Fig. S5.**
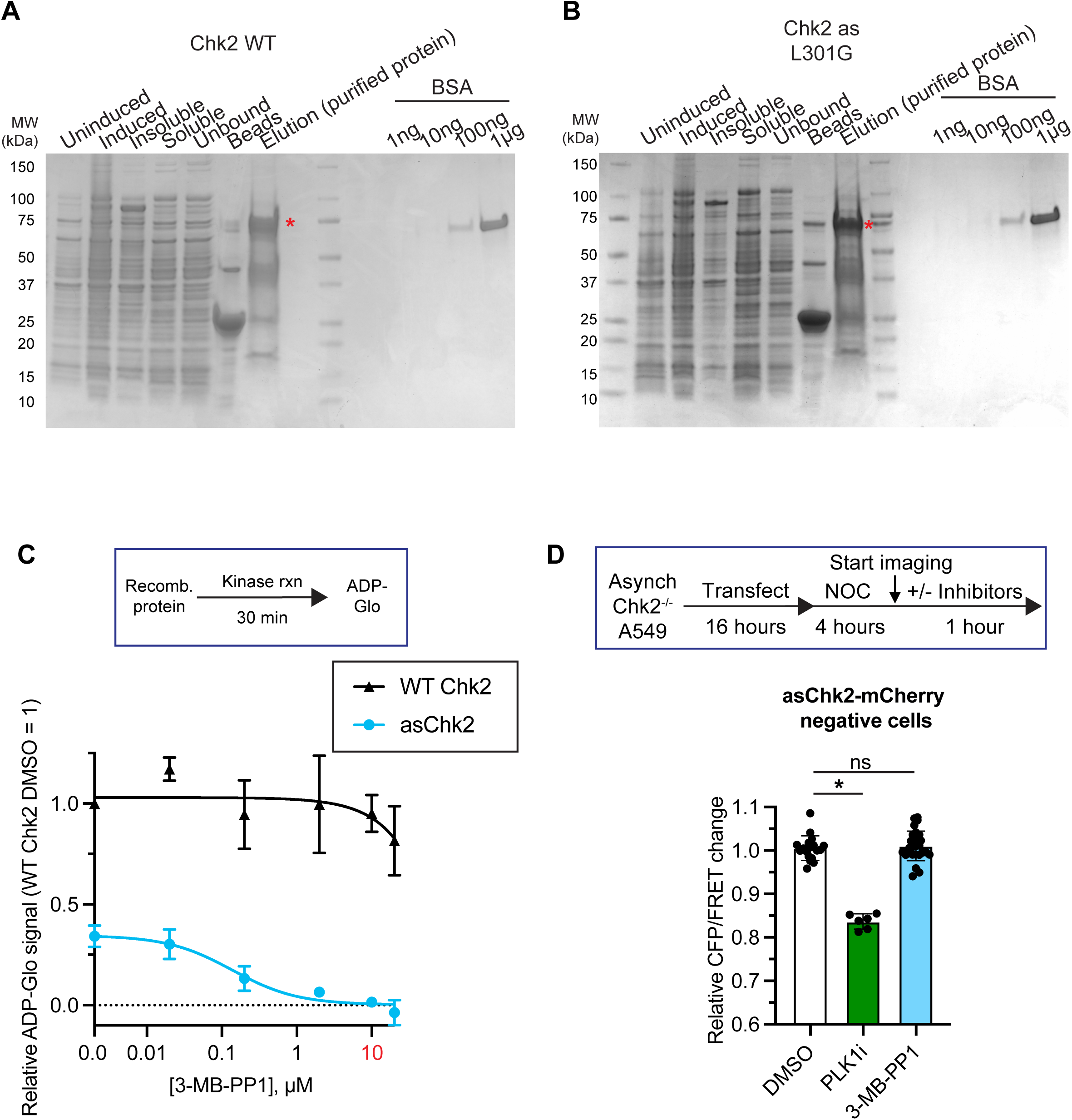
(A-B) GelCode™ Blue Safe Protein Stain of SDS-PAGE gels of recombinant Chk2-WT (A) or asChk2 (B) purification and intermediates. Red * indicates full-length Chk2. (C) ADP-Glo quantification of kinase activity in the presence of DMSO or varying concentrations of 3-MB-PP1. Error bars SEM of ≥3 experimental replicates. Values are normalized to WT Chk2 DMSO-treated kinase activity. Data is the same as in Fig. 1F (D) Top, experimental setup as in Fig. 1G. Bottom, quantification of relative change in CFP/FRET 1 hour after addition of inhibitors or DMSO in cells that are not expressing asChk2-mCherry. Each point represents one cell normalized to the average value in the DMSO condition for each replicate. Error bars SD of individual cell values, >6 cells per condition.

**Fig. S6.**
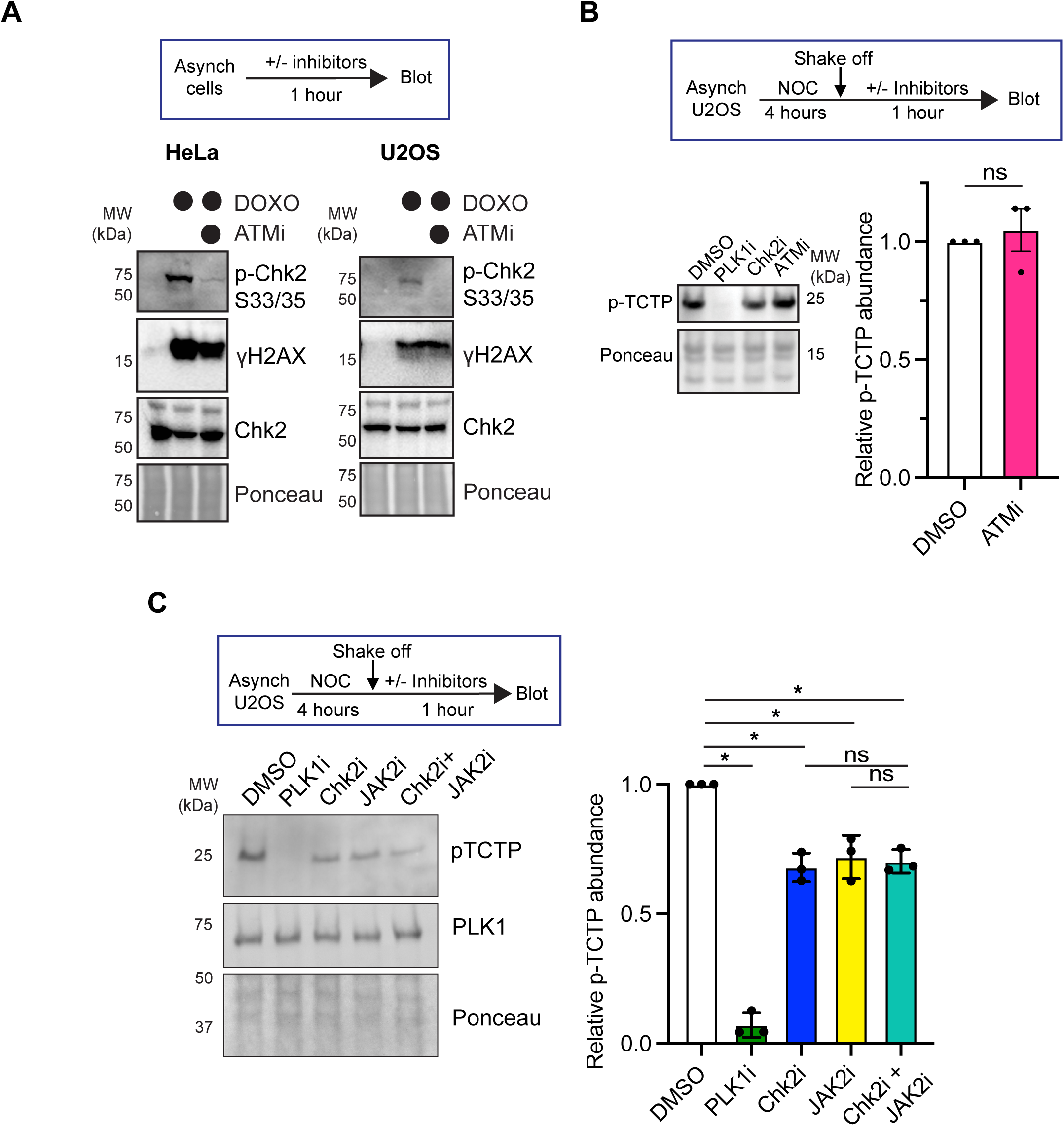
(A) Representative western blots of HeLa and U2OS cells untreated, treated with doxorubicin (DOXO, 2μM), or DOXO plus ATMi for 1 hour. (B) Quantification of p-TCTP abundance in mitotic U2OS cells treated for 1 hour with PLK1i (1μM BI-2536), Chk2i (10μM BML-277), ATMi (10μM KU55933), or equal volume DMSO as a vehicle control. Left, representative western blot. Right, quantification of 3 independent experiments. Western blot image for p-TCTP is the duplicate of U2OS image from Fig. S1A. ns, not significant, two-tailed t-test. (C) U2OS cells collected and treated as in Fig. 2C. Isolated mitotic cells were treated for 1 hour with PLK1i (1μM BI-2536), Chk2i (10μM BML-277), JAK2i (5μM JAK2 inhibitor IV), combination of Chk2i and JAK2i, or equal volume DMSO as a vehicle control. Following inhibitor treatment, cells were collected and immunoblotted. Left, representative western blot. Right, quantification of 3 experimental replicates. Each point demarcates value from biological replicate. Error bars SD. *p<0.05, one-way anova with Bonferroni multiple comparison correction. ns, not significant.

**Fig. S7.**
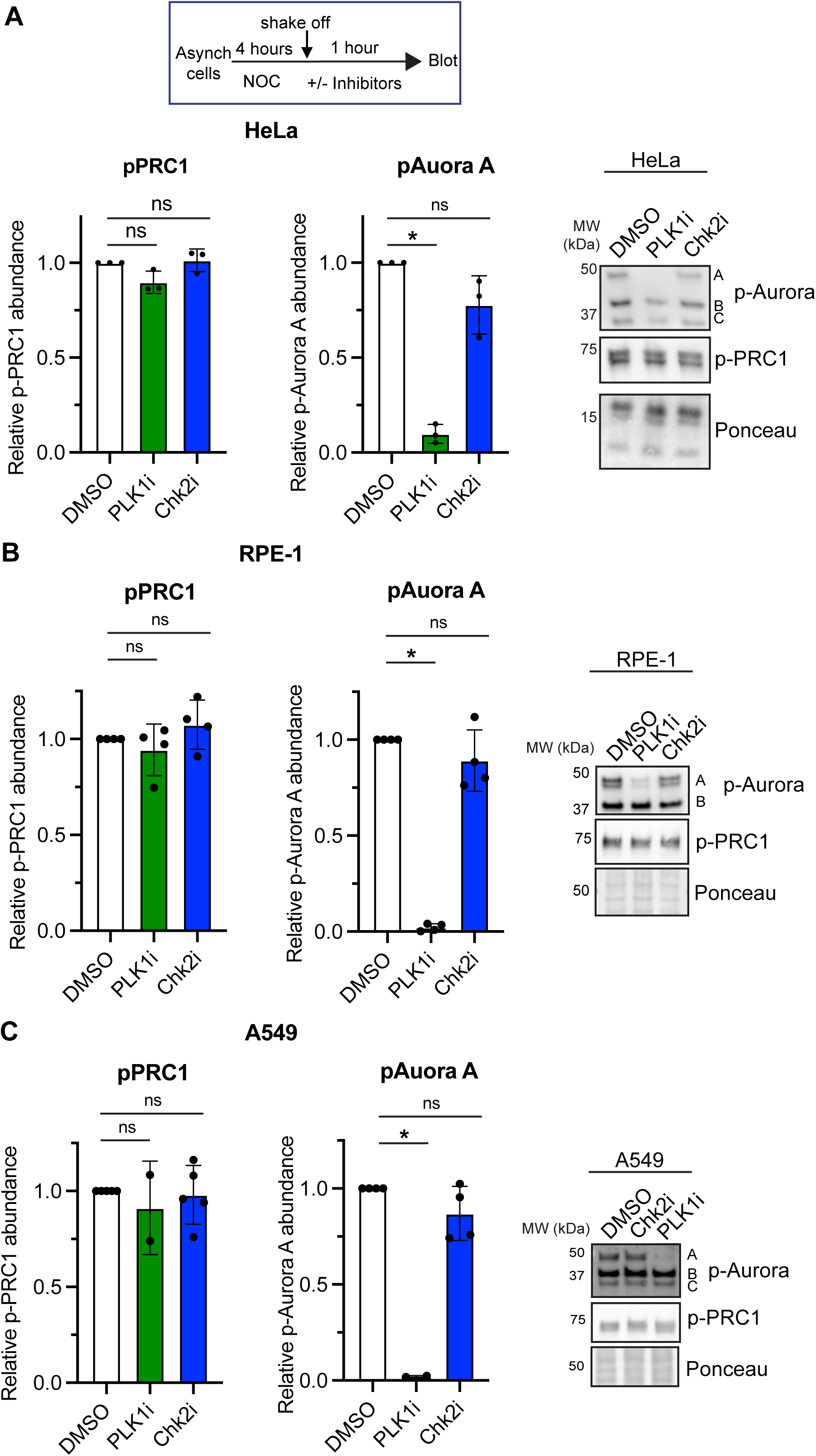
(A-C) Quantification (left) and representative western blots (right) of phospho-Aurora A T288 and phospho-PRC1 T481 in HeLa (A), RPE-1 (B), and A549 (C) cells treated as in Figure 1A. The ponceau image for the A549 cells is from the same experiments as in Fig. S6C. Ponceau imafe from HeLa cells is the same image as in Fig. S1A. *p<0.05, two-tailed t-test. ns, not significant (p>0.05).

**Fig. S8.**
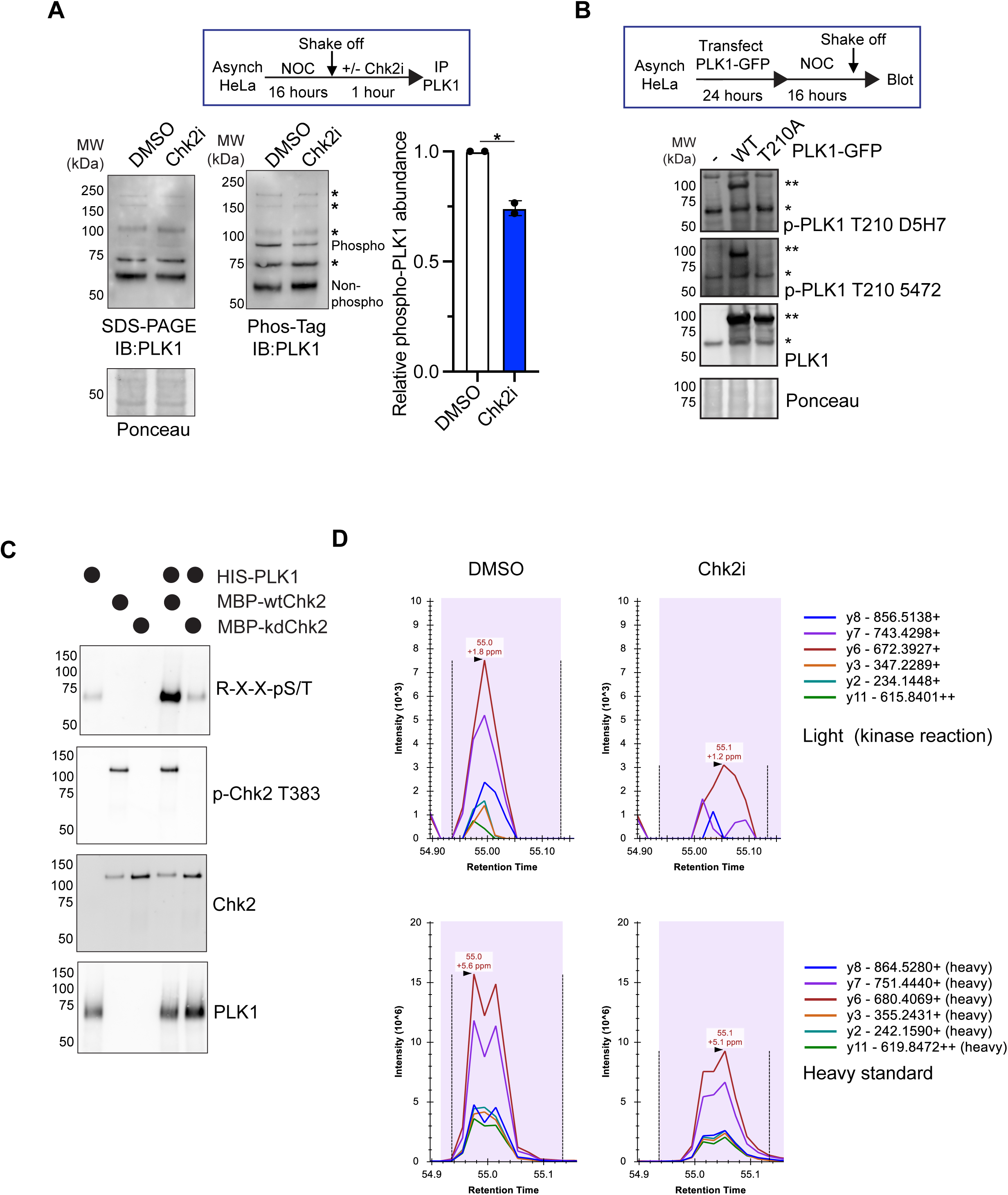
(A) Left, immunoblot of mitotic HeLa cells treated with a vehicle control or Chk2 inhibitor for 1 hour. Right, immunoblot of the same samples on a PhosTag gel. The putative phosphorylated specifies of PLK1 is indicated. *, uncharacterized bands recognized by PLK1 antibody. Right, quantification of phosphorylated PLK1 (slower-migrating PLK1 band on PhosTag gel) normalized to the total PLK1 protein abundance (68kDa band on SDS-Page gel). Each point represents one biological replicate. (B) Immunoblotting of mitotic HeLa extracts either untransfected (-) or transfected with GFP-PLK1 WT (T210) or T210A. *, endogenous PLK1 protein; **, exogenous (T210wt or T210A) PLK1 protein. (C) *In vitro* kinase assay with HIS-PLK1, MBP-wild-type(wt)Chk2, and/or kinase-dead T383A T387A (KD) MBP-Chk2. phospho-Chk2 T383 signal is only present in conditions with active Chk2. Kinase reaction performed as in Fig. 3D. (D) Phosphorylation site validation by targeted PRM mass spectrometry method. The unique MS2 ions of high resolution (different colored traces) were manually inspected with Skyline visualization. The synthetic lysine-labeled heavy phosphopeptide of (phos)TLC(Carbamidomethylation)GTPNYIAPEVLSK(+8Da) was used as a reference for relative quantification. Upper panel: the MS2 ions of light phosphopeptide (phos)TLC(Carbamidomethylation)GTPNYIAPEVLSK quantified and visualized by Skyline from DMSO (left) and Chki2 (right) sample. Lower panel: the MS2 ions of lysine-labeled heavy phosphopeptide (phos)TLC(Carbamidomethylation)GTPNYIAPEVLSK(+8Da) quantified and visualized by Skyline from DMSO (left) and Chki2 (right) sample.

**Fig. S9.**
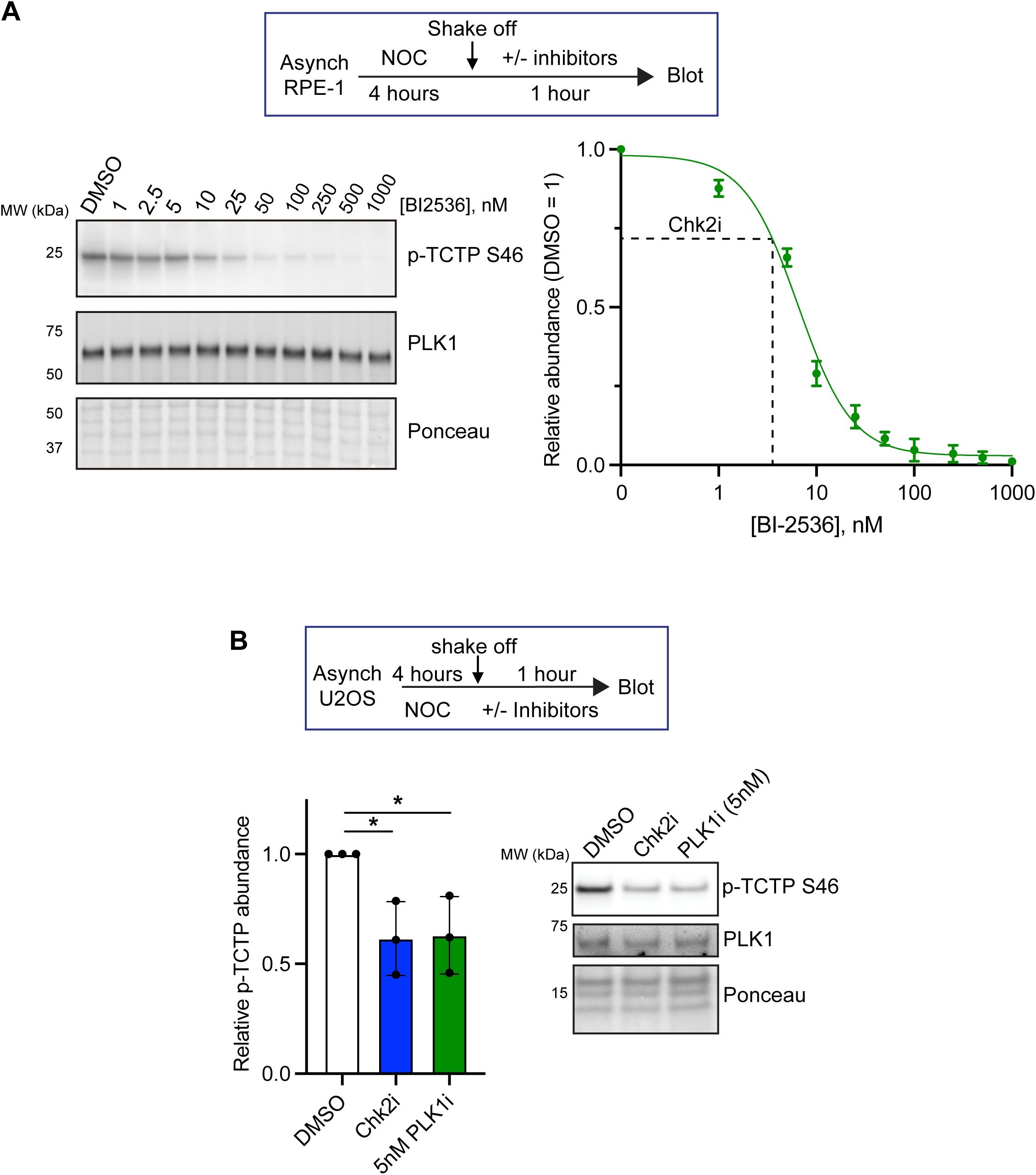
(A) Top, experimental setup. Representative western blot (left) and quantification (right) of nocodazole-arrested RPE-1 cells treated for 1 hour with varying concentrations of PLK1i BI-2536 as marked or a DMSO vehicle control. Error bars SEM of 3 experimental replicates. Dashed line marks the value of the average change in p-TCTP following Chk2i from Figure 1A. (B) Top, experimental setup. Representative western blot (left) and quantification (right) of nocodazole-arrested U2OS cells treated for 1 hour with 5nM PLK1i BI-2536, Chk2i (10μM BLM-277), or a DMSO vehicle control. Error bars SD of 3 experimental replicates. *p<0.05, two-tailed t-test.

**Fig. S10.**
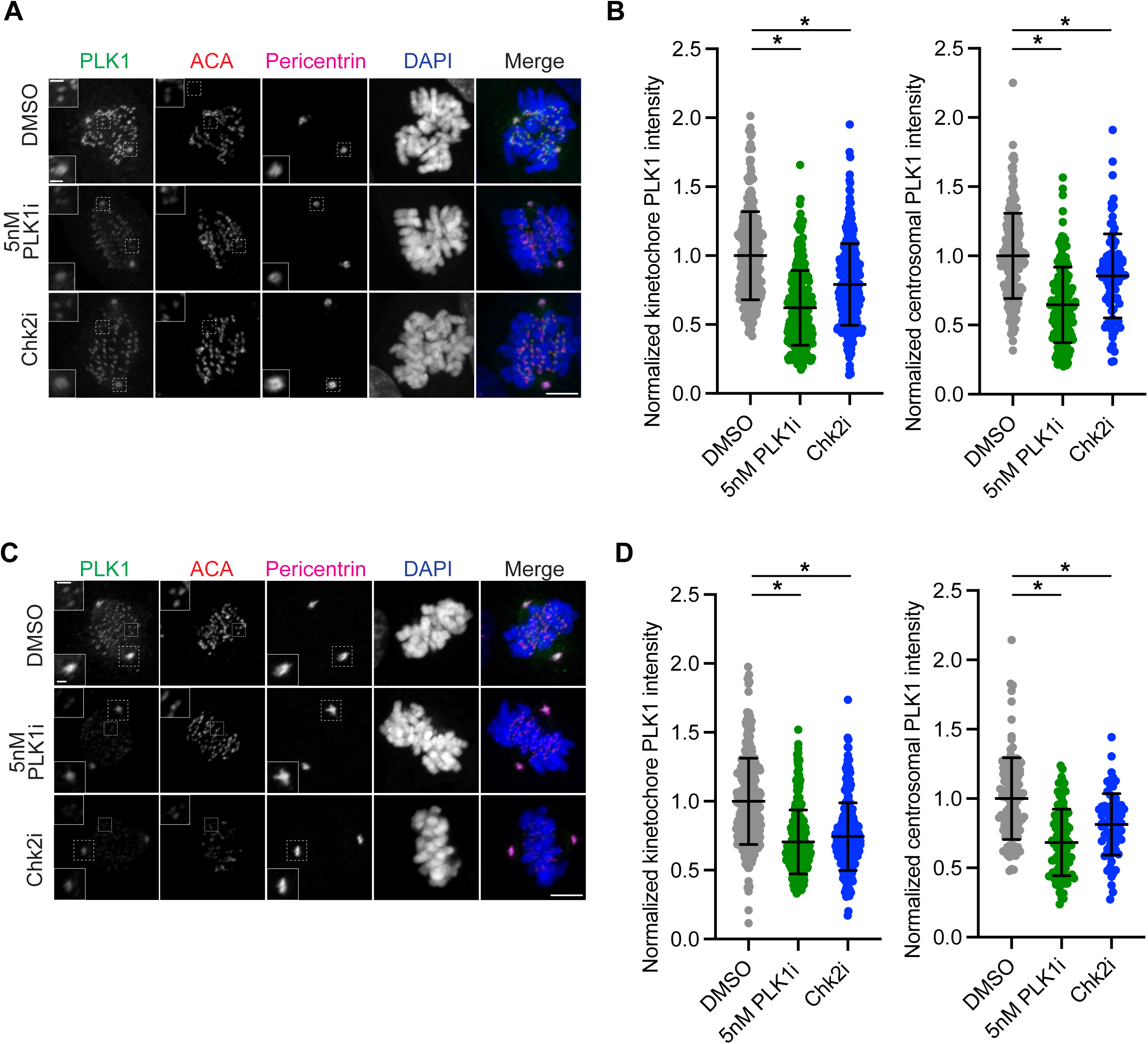
(A) Representative z-projected image of asynchronous prometaphase RPE-1 cells treated for 1 hour with 5nM PLK1i (BI-2536), Chk2i (10μM BLM-277), or a DMSO vehicle control. Cells were stained with DAPI and antibodies against PLK1, centromeres (anti-centromere antibody, ACA), and pericentrin, a centrosome component. Scale bar on large panels, 5μm. Scale bar on insets, 1μm. (B) Quantification of kinetochore-associated (left) or centrosome-associated (right) PLK1 staining in cells treated as in (A). Quantification was performed on a single in-focus z-slice. Points are individual kinetochores (left) or centrosomes (right). 1 centrosome and 4-10 in-focus kinetochores were analyzed per cell. Data are pooled from 4 independent experiments after normalization of each replicate to the mean DMSO value, ≥95 cells total. *p<0.05, two-tailed unpaired t-test of replicate mean values. (C-D) As in (A-B), analysis was performed on asynchronous metaphase cells. Each point represents one kinetochore (left) or centrosome (right) value. Data are pooled from 3 experimental replicates, ≥74 cells total.

**Fig. S11.**
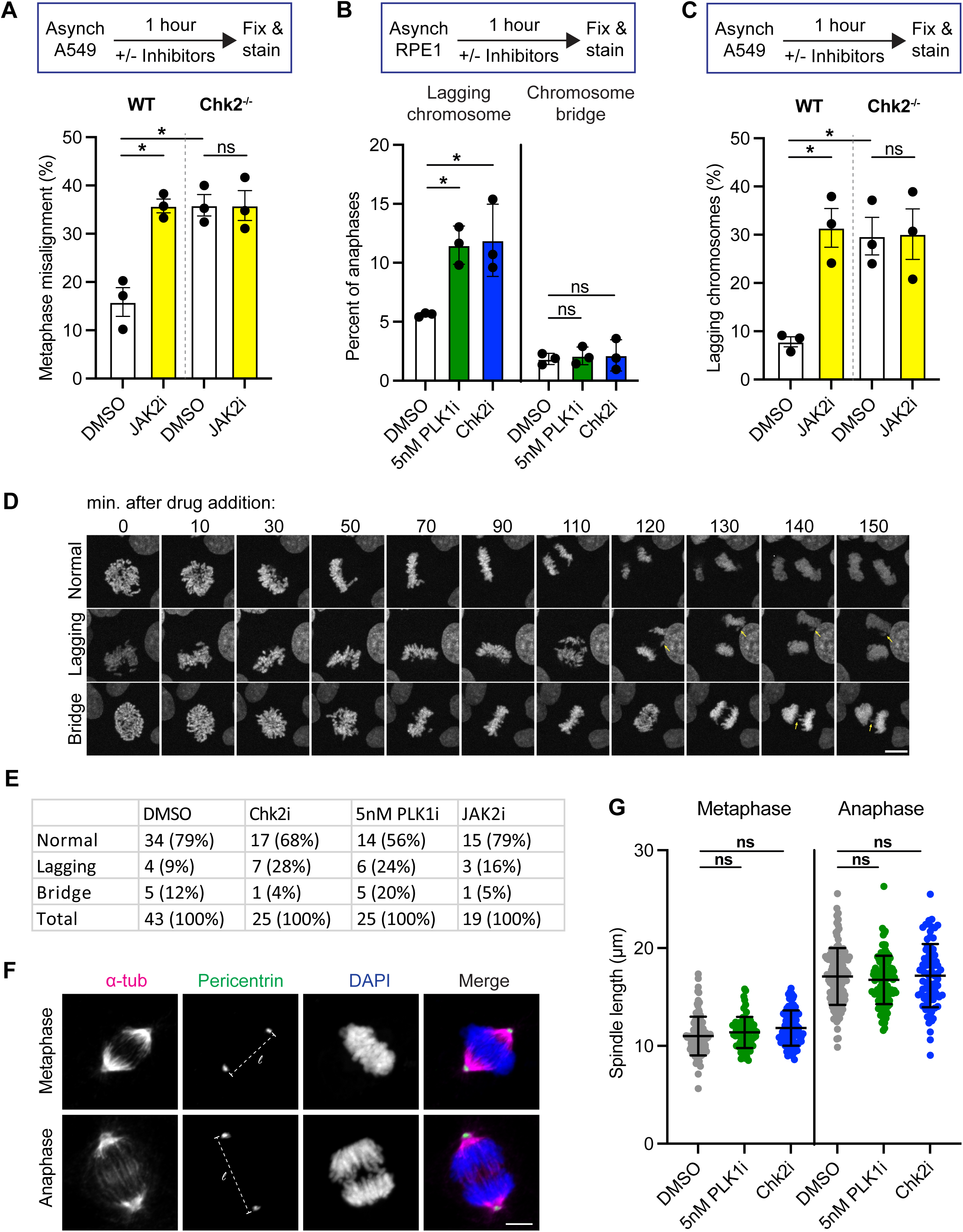
(A) Quantification of metaphase chromosome misalignment in A549 WT or Chk2^-/-^ cells treated for 1 hour with JAK2i or DMSO. Each point represents the rate from an experimental replicate (≥146 metaphases total per condition, ≥38 metaphases per condition per replicate). *p<0.05, unpaired one-way ANOVA with Bonferroni multiple comparison correction. ns, not significant. (B) Quantification of mitotic defects in RPE-1 cells treated for 1 hour with 5nM BI-2536 (PLK1i), Chk2i (10μM BLM-277), or a DMSO vehicle control. Each point represents the rate from an experimental replicate (≥240 anaphases total per condition, ≥52 anaphases per condition per replicate). *p<0.05, two-tailed t-test of replicate averages. ns, not significant (p>0.05). (C) Quantification of lagging chromosomes in A549 WT or Chk2^-/-^ cells treated for 1 hour with JAK2i or DMSO. Each point represents the rate from an experimental replicate (≥94 anaphases total per condition, ≥27 anaphases per condition per replicate). *p<0.05, unpaired one-way ANOVA with Bonferroni multiple comparison correction. ns, not significant. (D) Representative still images of quantified anaphase defects from live-cell imaging of asynchronous HeLa H2B-GFP cells. Asynchronous cells in prophase were imaged (time 0) and then the media was replaced with fresh media containing DMSO, 5nM PLK1i, Chk2i, or JAK2i. Numbers above images denote the time after drug addition in minutes. Yellow arrows are pointing out lagging chromosome (middle panel) and bridge (bottom panel). Images are maximum intensity z-projections. Scale bar, 10μm. (E) Quantification of chromosome segregation defects in HeLa H2B-GFP cells. Numbers denote the number of anaphases observed per phenotype and the number in parentheses denotes the percentage of observed anaphases per phenotype per condition. The numbers of cells reported are combined from two independent experimental replicates. (F) Representative z-projected image of asynchronous metaphase (top) or anaphase (bottom) RPE-1 cells. Cells were stained with DAPI and antibodies against pericentrin, a centrosome component, and alpha tubulin. Scale bar, 5μm. l indicates spindle length measurement in (G). (G) Quantification of spindle length (measured by intracentrosomal distance) in asynchronous RPE-1 cells treated as in (A). Each point represents one cell combined from ≥3 experimental replicates, ≥69 cells total per condition. ns, not significant, two-tailed t-test of replicate averages.

**Fig. S12.**
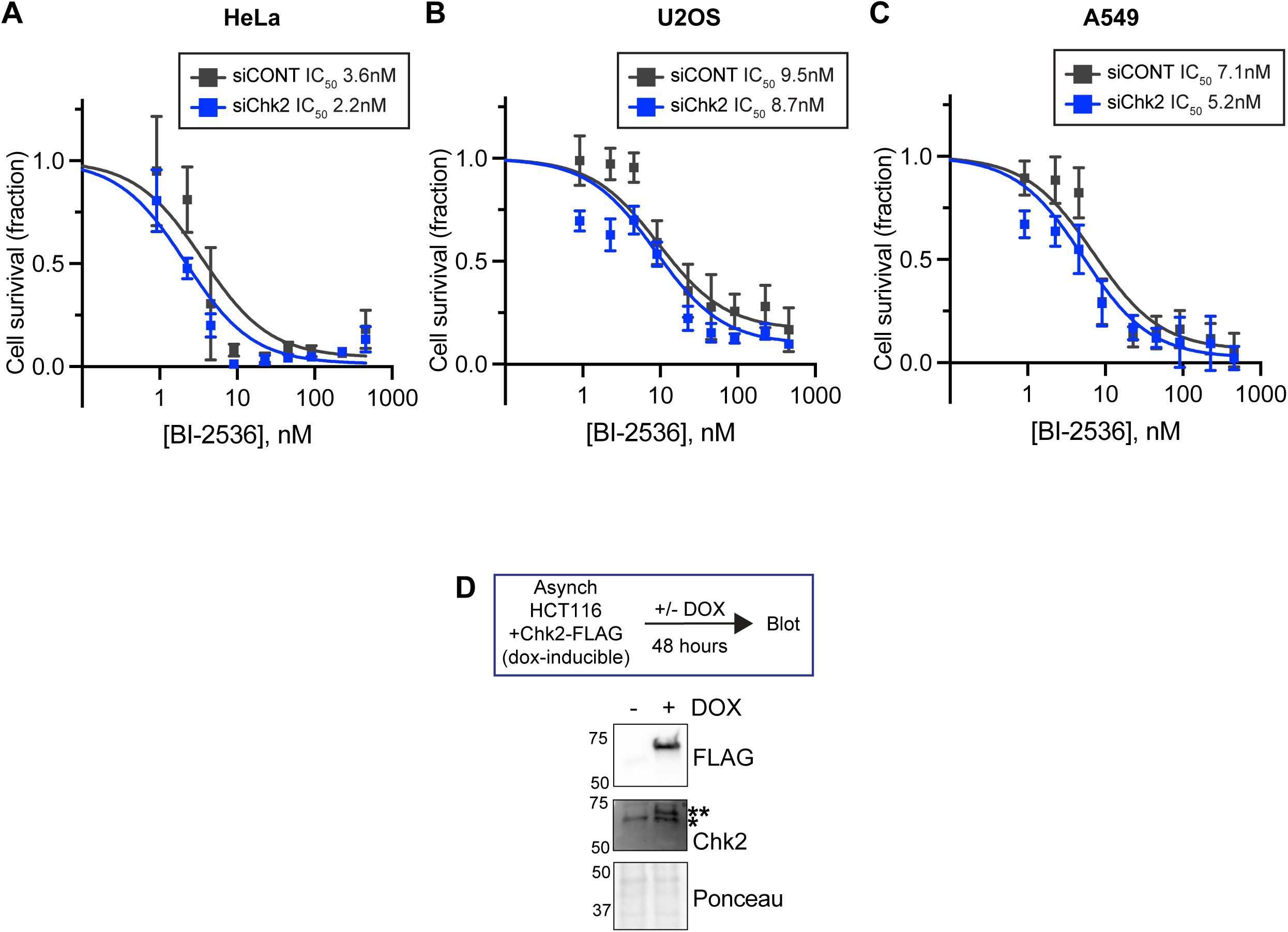
(A-C) Relative cell viability (DMSO-treated cells = 1) of HeLa (A), U2OS (B), and A549 (C) cells following 3-day siRNA treatment and treatment with PLK1 inhibitor BI-2536. Error bars SEM of ≥3 replicate averages. Values in Fig. 5C are derived from this data. (D) Western blot showing relative Chk2 expression in asynchronous cells following 48hr induction of Chk2-FLAG expression. *Endogenous Chk2, **3X-FLAG-Chk2

## Materials and Methods

### Cell lines and culture medium

U2OS (ATCC, HTB-96), RPE-1 hTERT (ATCC, CRL-4000), RPE-1 hTERT asPLK1-GFP (gift from Burkard lab), HeLa (Abcam, ab255928), PANC-1 (ATCC, CRL-1469), HeLa H2B-GFP (Sigma-Aldrich scc117), and 293T (ATCC, CRL-3216) cells were cultured in Dulbecco’s modified Eagle’s medium (DMEM; Gibco) supplemented with 10% fetal bovine serum (FBS; Gibco) and 1% penicillin/streptomycin (P/S) (Gibco). HCT-116 (ATCC, CRL-247) cells were cultured in McCoy’s 5A medium (ATCC 30-2007) supplemented with 10% FBS and 1% P/S. A549 WT (Abcam, ab275463) and Chk2 knockout cells (Abcam, ab276098) were grown in Ham’s F-12K Kaighn’s Modification (Gibco 21127-022) supplemented with 10% FBS and 1% P/S. All cells were cultured at 37°C with 5% CO_2_. HCT-116 cells stably expressing the Chk2 rescue construct were cultured in the presence of 0.5mg/ml Geneticin (G418, Gibco 10131-027). 48 hours prior to drug sensitivity assays, cells were grown in media without selection agents.

### Cell transfection

All cells were transfected with plasmids using the Lipofectamine™ 3000 Transfection Reagent (Invitrogen) according to manufacturer instructions. siRNA experiments were performed by transfection of either the *Silencer*™ Select Negative Control No. 1 siRNA (ThermoFisher, item number 4390843) or *Silencer*™ Select Pre-designed Chk2 siRNA (ThermoFisher, item number 119278) using the Lipofectamine™ RNAiMAX Transfection Reagent (Invitrogen) according to manufacturer instructions. 1pmol of siRNA was transfected per 96 well for viability experiments and 25pmol of siRNA was transfected per 6 well. plk1 T210A was a gift from Michael Yaffe (Addgene plasmid # 132965; http://n2t.net/addgene:132965; RRID:Addgene_132965). plk1 WT GFP-myc was a gift from Michael Yaffe (Addgene plasmid # 132963; http://n2t.net/addgene:132963; RRID:Addgene_132963).

### Viral Infection

Doxycycline-inducible 3X-FLAG Chk2 lentivirus was generated via co-transfection of 293T cells with pSLIK_neo_3xflag_CHEK2, which was a gift from Kevin Janes (Addgene plasmid #136536; http://n2t.net/addgene:136536; RRID:Addgene_136536), pCMV dR8.2 dvpr (Addgene 8455, gift from Bob Weinberg), and pCMV-VSV-G (Addgene 8454, gift from Bob Weinberg) using Lipofectamine™ 3000 Transfection Reagent (Invitrogen) according to manufacturer instructions. Viral supernatant was harvested 3 days after transfection and added to HCT-116 cells cultured in McCoy’s 5A supplemented with 1% P/S, 10% FBS, and 8µg/mL polybrene infection/transfection reagent (Sigma-Aldrich). HCT-116 cells were selected and grown in the presence of 0.5mg/ ml Geneticin (G418, Gibco 10131-027).

### Western blotting

Cells were collected for blotting by either collecting non-adherent shaken-off mitotic cells or by trypsinization (interphase cells). Cells were spun down, washed 1X with PBS, and then resuspended in 2x SDS Sample Buffer (0.1M Tris-HCl, 4% sodium dodecyl sulfate, 3mM Bromophenol blue, 2M glycerol, 20% beta-mercaptoethanol). The protein samples were lysed with an insulin syringe and boiled at 95°C for 15 min before being run on an SDS-Page gel. 50μM PhosTag acrylamide (FUJIFILM Wako Chemicals U.S.A. Corporation) within hand-cast 7.5% acrylamide gels was used to separate phosphorylated PLK1 (Fig. S8A). Protein gels were transferred to 0.45 μM nitrocellulose membranes using the Trans-Blot Turbo Transfer System (Bio-Rad). Membranes were stained with Ponceau S staining solution (0.5% (w/v) Ponceau S, 1% glacial acetic acid) to visualize total protein loading, imaged, and then Ponceau was removed by washing with 1X Tris-Buffered Saline + 0.1% Tween-20 (TBS-T). Membranes were blocked with TBS-T + 5% bovine serum albumin (BSA) or 5% milk for at least 1h at room temperature with gentle rocking. Primary antibodies were diluted in blocking solution, then added to membranes to incubate overnight at 4°C with gentle agitation. Membranes were washed 3X with TBS-T for 5 min before incubation with secondary antibody. Secondary antibodies conjugated to with fluorophores or horseradish peroxidase (HRP) were diluted 1:5000 in 1% BSA and added to membranes for at least 1 hour at room temperature, followed by 2 additional washes to remove unbound secondary antibody. Membranes were incubated with Clarity Western ECL substrate (for HRP-conjugated antibodies; Bio-Rad) and imaged on the ChemiDoc MP Imaging system (Bio-Rad).

### Immunofluorescence

Cells were grown on coverslips and treated with the indicated drugs, then fixed with 3.5% paraformaldehyde for 15 min, permeabilized with Phosphate Buffered Saline (PBS) + 1% Triton X-100 for 15 min and blocked with PBS +1% BSA for 10 min. Coverslips were then incubated with primary antibody diluted in PBS +1% BSA at 4°C overnight. Coverslips were then washed with PBS to remove unbound primary antibody. Secondary antibodies conjugated to fluorophores and 1mg/mL DAPI were diluted 1:1000 in PBS +1% BSA before being added to coverslips for 1hr at room temperature. Cells were washed again with PBS and mounted onto glass slides using ProLong Gold Antifade Mountant (Invitrogen) and left to dry overnight. All images were acquired on a Nikon ECLIPSE Ti2 W2 spinning disk confocal microscope with a Hamamatsu Fusion sCMOS camera. Image acquisition was managed using NIS-Elements (Nikon).

### Fluorescence and western blot quantification

All images were analyzed in Fiji. For western blot quantification, regions of interest were drawn around the band and the area and mean gray value for each band was recorded. A mean gray value for the background in a negative space immediately adjacent to the band was also recorded. The value for each band was calculated by subtracting the mean gray value for the background from the mean gray value for the band and then multiplying that background-subtracted value by the band’s area. All values were then normalized to similarly background-subtracted values in Ponceau S staining (unless otherwise indicated), normalized to the control, and graphed. Similarly, immunofluorescence values were calculated by measuring the fluorescence intensity for an area of interest and then subtracting out a mean gray value picked from a region without cells. For p-TCTP S46 immunofluorescence, a no primary control coverslip was imaged alongside experimental samples, and the average gray value from >5 mitotic cells on the no primary control was the subtracted background value for experimental samples.

### FRET

Plk1 FRET sensor c-jun substrate was a gift from Michael Lampson (Addgene plasmid #45203; http://n2t.net/addgene:45203; RRID:Addgene 45203). Cells were transiently transfected with a and plated onto glass-bottom plates. The next day, cells were arrested in nocodazole for 4 hours prior to imaging. Live-cell imaging was performed with a heated stage at 37°C and 5% CO_2_. CFP, YFP, FRET, and brightfield images were taken for each cell prior to addition of drugs. After initial image acquisition, the media for each condition was replaced with fresh media containing nocodazole and DMSO or a small-molecule inhibitor against the kinase of interest. Imaging was resumed for 1 hour after addition of drugs. CFP/FRET ratios were calculated by subtracting background values for FRET and CFP at both the initial timepoint (T_0_, before addition of inhibitors) and the final timepoint (T_60_, 1 hour) and calculating (CFPt_60_/ FRETt_60_) / (CFPt_0_/ FRETt_0_). Ratio values were then normalized so that the mean CFP/FRET ratio change after 1 hour for the DMSO condition equals 1. Replicates were then combined and plotted as SEM of replicate averages (Fig. 1D) or as individual cell values (Fig. 1G, S4C, S5D).

#### Live-cell imaging

Asynchronous HeLa cells stably expressing H2B-GFP were cultured in a 24-well glass-bottom dish (Cellvis). The dish was mounted on a Tokai Hit STX stage-top incubator (Spectra Services) to maintain an imaging environment of 37°C with 5% CO2. Z-stacks were taken at 60x for each cell with a range of 8µm at 1µm steps. After identifying prophase cells, initial images of the cell in DIC and 488 image were taken, and then the media was replaced with fresh media containing 10µM Chk2i, 5nM BI-2536, 5uM JAK2i, or an equal volume DMSO control. The remaining images were taken at 5-minute intervals over the course of 2.5 hours.

#### PLK1 Immunoprecipitation

Following nocodazole arrest and inhibitor treatment, U2OS cells were washed with ice-cold dPBS (Gibco) and lysed in lysis buffer (20mM HEPES [pH 7.4], 150mM KCl, 10% glycerol, 5mM DTT, 1% Triton X-100, 1 Pierce Protease Inhibitor Tablet [EDTA-free], and 1 phosphatase inhibitor PhosSTOP™ Roche tablet). Crude cell extract was lysed with an insulin syringe and pre-cleared by incubation with 3 μg Normal Rabbit IgG (Cell Signaling Technology) and 10uL Pierce™ Protein A/G Magnetic Beads (Thermo Scientific 88803) with constant rotation at 4°C for 4 hours. The beads were removed with centrifugation at 23000 rcf for 1 minute. The remaining pre-cleared cell extract was incubated overnight with 5 μg of Rabbit Anti-PLK1 antibody (Millipore) with constant rotation at 4°C. 25uL of Protein A/G Magnetic Beads were added to the suspension and samples were incubated for an additional 4 hours at 4°C with constant rotation. The beads were washed 5 times with Lysis Buffer and vortexing and were then eluted SDS elution buffer (50 mM Tris-HCl [pH 8.0], 1 mM EDTA [pH 8.0], 1% SDS). Immunoprecipitated protein was eluted at 95°C for 10 minutes. Samples were diluted with 2x Sample Buffer and prepared for western blotting as described above.

### Chk2 purification

Wild-type and analog-sensitive Chk2 ORF sequences were cloned in the pGex6p-1 plasmid (Genscript, see plasmid construction section for details) and transformed into BL21 bacterial cells (New England Biolabs). Protein expression was induced by growing a dense overnight culture, diluting to an OD_600_ value of 0.1, growing the culture until OD_600_ equals 0.4, and then adding 1mM IPTG (Bio Basic item number 367-93-1) for 4 hours at 37°C with shaking. Cells were collected by centrifugation and frozen at -80°C. Cells were resuspended in 10mLs chilled lysis buffer (50mM tris PH 7.4, 150mM NaCl, 1mM DTT, 50mM NaF, 5mM EDTA, 1% glycerol supplemented with protease inhibitor tablet (Pierce), pH adjusted to 7.6) and sonicated to lyse. The lysate was then centrifuged at 45,000xg for 20 minutes at 4°C, and the soluble fraction was collected for Chk2 affinity purification. Chk2 was immunoprecipitated from the soluble fraction by addition of Glutathione-conjugated magnetic agarose beads (Pierce 78601) overnight with gentle rocking. After thorough washing in ice cold cleavage buffer (50mM tris pH 7, 150mM NaCl, 1mM DTT), Chk2 was eluted from the beads by addition of PreScission Protease (Genscript Z02799) overnight rocking at 4°C. Finally, the protease was trapped on the beads by addition of additional GST conjugated beads and 1 hour incubation with rocking at 4°C. The GST beads were isolated with a magnet, and purified Chk2 was collected in the supernatant and combined 1:1 with freezing buffer (100mM tris, 300mM NaCl, 10mM DTT, 82% glycerol) and stored at -80°C in single-use aliquots.

### ADP-Glo™

ADP-Glo™ (Promega) was performed according to manufacturer instructions. Briefly, kinase reactions were performed with 10-100ng of the kinase of interest (optimized for each kinase) and 0.5ug Chktide substrate (derived from human CDC25C protein isoform A amino acids 205– 225, used in Chk2 kinase reactions) or Casein, PLK1 and Aurora A sustrate, in manufacturer kinase reaction buffer (40mM tris pH 7.5, 20mM MgCl2, 0.1 mg/mL BSA, 1mM DTT, and 100μM ATP) at 30°C for 30 minutes with shaking with indicated concentrations of inhibitor or DMSO control. A control reaction with no enzyme was also included. The reaction was then cooled to room temperature and incubated with equal volume of ADP Glo reagent for 40 minutes before then being incubated with kinase detection reagent for 1 hour. Reactions were transferred to a 384 well plate and luminescence was recorded on a BioTek Synergy H1 microplate reader with an integration time of 0.5ms. Values recorded for the substrate-only reaction were subtracted from experimental values, and these subtracted experimental values were normalized to vehicle control (DMSO) reactions. Reactions were performed in a minimum of three experimental replicates.

### *In vitro* kinase assays & recombinant proteins

GST-Chk2 used in Fig. S3A and S8D was purchased from Promega (V4020). HIS-PLK1 used in ADP-Glo was from Promega (V2841). HIS-PLK1 used in Fig. 3D and S8D *in vitro* reactions was from Abcam (AB51426-1002). HIS-Aurora A was purchased from Abcam (AB42595). Wild-type or T383A T387A kinase-dead MBP-Chk2-6xHIS was a gift from Jesse Reinhart lab ^66^. Details of individual *in vitro* kinase reactions are described in the corresponding figure captions. Curve fit for *in vitro* kinase assay inhibitor IC_50_ calculations were calculated with nonlinear regression with standard slope, equal to a Hill slope of -1.0. All kinase reactions were performed in kinase assay buffer (20mM 3-morpholinopropane-1-sulphonic acid (MOPS), pH 7.2, 25mM MgCl_2_, 500μM ATP, 1 mM DTT) at 30°C with gentle agitation. Following incubation, reactions were quenched by adding EDTA to a final concentration of 5mM before being analyzed by immunoblotting. Reactions to be analyzed by mass spectrometry were flash-frozen in liquid nitrogen.

### Parallel Reaction Monitoring (PRM) Quantification of the Phosphorylation Site

The Orbitrap Fusion Lumos Tribrid based mass spectrometer MS (Thermo Fisher Scientific) was used for the targeted proteomic measurement using the Parallel Reaction Monitoring (PRM) mode ^85^. And Skyline ^86^ was used to visualize and report the results. Briefly, gel bands were processed by following a standard protocol ^87^ with trypsin (Promega) digestion at a 10-ng/μL concentration overnight. After C18 purification, the digest peptides from DMSO and Chk2i conditions were respectively spiked with the synthetic lysine-labelled heavy phospho-peptide of (phos)TLC(Carbamidomethylation)GTPNYIAPEVLSK(+8Da) that was synthesized by Thermo Fisher Scientific. The resulted samples were injected to MS in which PRM measurement was performed for confirming the endogenous phospho-peptide identity and for the relative quantification, using 500fmol of the heavy phospho-peptide as the reference. The theoretical peptide precursor m/z values of the light and heavy versions used in PRM analysis were generated by Skyline. The isolation window was set to be 1.4m/z. The Orbitrap resolution for PRM was set at 30,000, AGC target 1.0e5, and maximum injection time 150 ms ^88^. A stepped higher energy collisional dissociation collision energy of 2% (centered at 28%) was used. The resultant PRM data were imported into Skyline for manual inspection and quantification.

### Plasmid construction

The plasmids generated in the study are Chk2 WT and analog-sensitive (L301G) in an N-terminal constitutive mCherry mammalian expression vector and pGEX-6p-1, an inducible bacterial expression vector. The open reading frame of wild-type Chk2 was subcloned by amplifying the open reading frame of Chk2 from pSLIK_neo_3xflag_CHEK2, which was a gift from Kevin Janes (Addgene plasmid #136536; http://n2t.net/addgene:136536; RRID:Addgene_136536). Constructs were validated with Sanger sequencing before use. The analog-sensitive L301G mutation in Chk2 was generated by site-directed mutagenesis where nonoverlapping primers (TTATATTGTTGGGGAATTGATGGAAGG, FW; TAATCTTCTGCATCAAAAAAG, RV) introducing the single-nucleotide change were used to amplify the mCherry-wtChk2 sequence, then were phosphorylated by polynucleotide kinase (PNK) and ligated together using T4 DNA ligase. Mutation was confirmed via sequencing (sequencing primer GGAAGTGGTGCCTGTGGAGAG).

### Cell line genotyping

DNA was extracted from HCT-116, A549 WT, and A549 Chk2^-/-^ cells using the Monarch^®^ Genomic DNA Purification Kit (NEB). Prior to Sanger sequencing, regions surrounding the putative mutation sites were amplified using Phusion High-Fidelity polymerase (NEB) with the below primers:

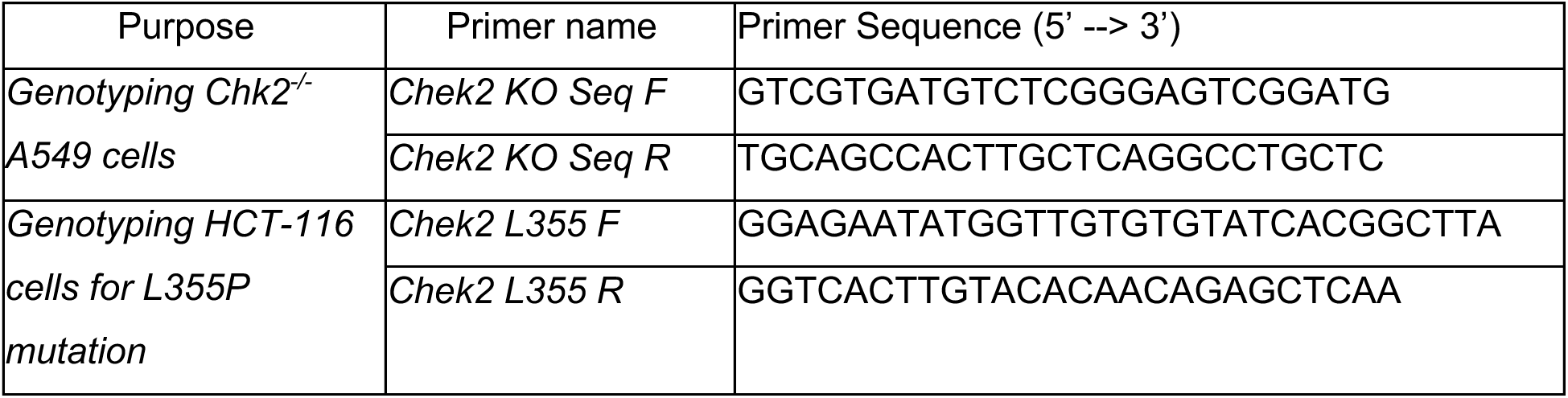

### Antibodies

Antibodies used for this study were: human anti-centromere antisera (ACA) (Antibodies Inc., 15-234-0001), alpha tubulin (Cell Signaling Technology, 3873), Chk2 (Cell Signaling Technology, 3440), cyclin B1 (Cell Signaling Technology, 12231), FLAG (Sigma-Aldrich F1804), GAPDH (Santa Cruz Biotechnology sc32233), phospho-Aurora A/B/C (Cell Signaling Technology D13A11), phospho-CDK1 Y15 (Fortis Life Sciences BLR101H), pericentrin (Abcam ab4448), PLK1 mouse (used in all western blots except for PhosTag, Millipore 05-844), PLK1 rabbit (used in Phos-tag gel and PLK1 IP, Millipore ABE2619), phospho-PRC1 T481 (Abcam ab62366), phospho-TCTP (Cell Signaling Technology 5251), R-X-X-phosphoS/T (Cell Signaling Technology 9614), TCTP (Cell Signaling Technology 5128), γH2AX (Novus NB100-74435), phospho-PLK1 T210 D5H7 (Cell Signaling Technology 9062), phospho-PLK1 T210 5472 (Cell Signaling Technology), phospho-Chk2 S33/35 (Cell Signaling Technology 2665), phospho-Chk2 T383 (Thermo Fisher Scientific PA5-37786)

### Inhibitors and small molecule reagents

Nocodazole (Cayman 13857100ng/mL), BI-2536 (Cayman 17388, concentration as indicated), BLM-277 (EMD Millipore Corp 220486, concentrations as indicated), 3-MB-PP1 (EMD Millipore Corp. 529582, concentrations as indicated), S-trityl-L-cysteine (STLC, Sigma Aldrich 164739, 10μM), Doxorubicin (DOXO, TCI D4193, 2μM), Etoposide (Etop, Cayman 12092, 10μM), MLN8237 (Aurora A inhibitor, Cayman 13602, 50nM), RO-3306 (CDK1 inhibitor, Cayman 15149, 5μM), MG132 (EMD Millipore Corp, 474790, 20μM), doxycycline (DOX, Alfa Aesar J67043), ATMi (KU55933, Cayman 16336, 10μM), DNA-PKi (M3814, Selleckchem S8586, 5μM), ATRi (AZ20, ChemCruz sc-503186, 10μM), JAK2i (JAK2 inhibitor IV, Thermoscientific j65506, 5μM).

### Drug sensitivity

Following plating in a 96-well plate, cells were treated in technical triplicates with inhibitors or an equal volume of DMSO for 3 days. Cells were washed thoroughly with PBS and then fixed and stained using crystal violet (0.5g per 100mL 20% methanol) for 1 minute. Cells were washed with 2x with water and crystal violet was measured using BioTek Synergy H1 microplate reader by measuring absorbance at 570 nm. For each replicate, technical triplicate values for each drug concentration were averaged. Data plotted represents the technical triplicate averaged value for each replicate. All data shown represent at least 3 independent biological replicates. Curve fit for cell viability IC_50_ calculations were calculated using nonlinear regression model with variable slope.

### Scoring mitotic defects

All mitotic defects were scored on fixed and stained coverslips. Lagging chromosomes were called when there was a DNA fragment (DAPI) that was also positive for ACA (anticentromere antibody) that was visibly isolated from the chromosome masses. DAPI signal that was ACA-negative was classified as acentric chromatin, and DAPI signal that was continuous between the segregating masses, regardless of ACA signal, was called as a chromosome bridge.

### Statistics and nonlinear regression analysis

For assays where individual points per cell are recorded, such as immunofluorescence and FRET, values from within a replicate were first tested for outliers using the ROUT test where Q = 1%. After removing outliers, values were normalized to the average value of the vehicle control. Unless otherwise indicated, statistics are performed on the average of the normalized replicate values using a two-tailed t-test. In places where multiple comparisons are performed, we used one-way ANOVA with Bonferroni’s correction.

